# Minimal interplay between explicit knowledge, dynamics of learning and temporal expectations in different, complex uni- and multisensory contexts

**DOI:** 10.1101/2021.03.06.434202

**Authors:** Felix Ball, Inga Spuerck, Toemme Noesselt

## Abstract

While temporal expectations (TE) generally improve reactions to temporally predictable events, it remains unknown how temporal rule learning and explicit knowledge about temporal rules contribute to performance improvements and whether any contributions generalise across modalities. Here, participants discriminated the frequency of diverging auditory, visual or audiovisual targets embedded in auditory, visual or audiovisual distractor sequences. Temporal regularities were manipulated run-wise (early vs. late target within sequence). Behavioural performance (accuracy, RT) plus measures from a computational learning model all suggest that temporal rule learning occurred but did not generalise across modalities, that dynamics of learning (size of TE effect across runs) and explicit knowledge have little to no effect on the strength of TE, and that explicit knowledge affects performance – if at all – in a context dependent manner: only under complex task regimes (unknown target modality) might it partially help to resolve response conflict while it is lowering performance in less complex environments..

## 1. Introduction

Gathering temporal information is an essential aspect of our life. Every day, we use temporal information as to when it is most likely to catch the bus, or in sports, we estimate when and where a ball has to be kicked, hit or caught. Computations resulting in temporal expectations and predictions have been studied in various different experimental paradigms and sensory systems (Nobre & Rohenkohl, 2014). There is converging evidence that temporal expectations (TE), at least in the younger population, improve performance for targets expected in time (Ball, Fuehrmann, Stratil, & Noesselt, 2018; Ball, Michels, Thiele, & Noesselt, 2018; Correa, Lupiáñez, & Tudela, 2005; Mathewson, Fabiani, Gratton, Beck, & Lleras, 2010; Nobre & Rohenkohl, 2014; Roach, Heron, & McGraw, 2006; Rohenkohl, Cravo, Wyart, & Nobre, 2012; Vatakis, Bayliss, Zampini, & Spence, 2007; Zanto et al., 2011). However, little is known about (1) how temporal rule learning develops over time (as reflected by learning onsets as well as the probability of correct responses on each trial), and (2) whether explicit knowledge about learned temporal regularities influences the strength of temporal expectations.

TE effects (faster and more accurate responses for targets expected in time) are traditionally determined by contrasting the average performance scores of expected and unexpected trials. However, this type of analyses ignores the fact that learning of certain features (here temporal rules) is a highly dynamic process. To capture the dynamics of learning processes, the use of state-spaced learning models (Smith & Brown, 2003; Smith et al., 2004) has been proposed. These models estimate a learning curve based on single-trial accuracy data which can be used to determine not only how fast an association or rule has been learned (i.e. to identify the first learning trial) but also how strongly the rule has been learned (Hargreaves, Mattfeld, Stark, & Suzuki, 2012; Smith et al., 2004). This type of modelling has been successfully applied to experiments with memory-association tasks, visuo-motor associative learning tasks, location-scene association and T-maze tasks (Clarke, Roberts, & Ranganath, 2018; Hargreaves et al., 2012; Smith et al., 2004; Wirth et al., 2003). However, to our knowledge the modelling of learning curves has not been applied to temporal rule learning so far, although it would allow for testing critical concepts about how temporal information is processed over time and whether temporal information is generalised across sensory systems. For instance, in multisensory paradigms (e.g. Alais & Burr, 2004; Ball, Fuehrmann, et al., 2018; Ball, Michels, et al., 2018; Driver & Noesselt, 2008; Noesselt et al., 2007, 2010; Parise, Spence, & Ernst, 2012; Starke, Ball, Heinze, & Noesselt, 2017; Werner & Noppeney, 2010) temporal rules might either be transferred between modalities or learned independently for each modality. Hence, depending on whether the data are best described by a single learning curve (information transfer) or modality-specific learning curves (individual rule learning) one could infer which learning form is the most likely for a given dataset/task. Similarly, temporal rules (e.g. the most likely target positions in a stimulus sequence) might be learned run-dependent (i.e. would be reset during runs, resulting in multiple learning curves; here only the local, run-wise context of expected and unexpected temporal positions determines the data) or there might be information transfer between runs (leading to one learning curve; after learning a temporal position in one run, participants successively shift their attention to the newly expected position in the following run). Additionally, learning curves can be utilised to identify the onset of learning and whether this onset differs across experimental conditions and tasks (Clarke et al., 2018; Hargreaves et al., 2012; Smith et al., 2004; Wirth et al., 2003).

Turning to the potential effects of explicit knowledge of temporal information, differences between implicit and explicit knowledge about temporal regularities are often studied by comparing rhythm with cueing paradigms: the processing of rhythms is usually assumed to be under bottom-up (implicit TE) control (de la Rosa, Sanabria, Capizzi, & Correa, 2012; Rohenkohl, Coull, & Nobre, 2011), while the processing and use of temporal cues is usually assumed to be under top-down (explicit TE) control (Coull & Nobre, 2008). This notion is in close resemblance to research on exogenous vs. endogenous orientation of spatial attention (Giordano, McElree, & Carrasco, 2009; Kurtz, Shapcott, Kaiser, Schmiedt, & Schmid, 2017; Müller & Rabbitt, 1989; Warner, Juola, & Koshino, 1990). Accordingly, several studies observed distinctions between ‘implicit’ and ‘explicit’ temporal expectations on the behavioural and neural level (Capizzi, Sanabria, & Correa, 2012; Correa, Cona, Arbula, Vallesi, & Bisiacchi, 2014; Coull, Frith, Büchel, & Nobre, 2000; Coull & Nobre, 2008; Mento, Tarantino, Sarlo, & Bisiacchi, 2013). Note that most previous studies do not directly assess participants’ explicit knowledge of the temporal manipulation or use different tasks and/or paradigms for the comparison of implicit and explicit TE. However, even under implicit experimental task regimes, participants might gain knowledge about the underlying temporal regularities via explicit rule extraction during statistical learning (Hannula & Greene, 2012; Henke, 2010; Turk-Browne, Scholl, Chun, & Johnson, 2009; Turk-Browne, Scholl, Johnson, & Chun, 2010) and perceptual learning (Seitz, 2017; Seitz & Watanabe, 2009). In turn, they may thus be able to utilize their explicit knowledge to solve a particular task (Taylor & Ivry, 2013). More importantly, any observed differences between implicit and explicit knowledge could also be driven by the differences in stimulation protocols such as the use of different stimulus material or effects of entrainment vs. no entrainment (Ball, Groth, Agostino, Porcu, & Noesselt, 2019; Correa et al., 2014). Thus, studies are needed that compare the effects of explicit knowledge within the same paradigm.

Previous non-TE focussed studies investigating the influence of explicit knowledge – within the same paradigm – have reported divergent effects: while very few have observed an increase in performance (as measured by accuracies or response times), others have found no or even detrimental effects, or changes in confidence rather than performance (Batterink, Reber, Neville, & Paller, 2015; Fairhurst, Travers, Hayward, & Deroy, 2018; Green & Flowers, 2003; Mancini et al., 2016; Preston & Gabrieli, 2008; Sanchez & Reber, 2013; Van den Bussche et al., 2013). Recently, we were able to demonstrate that visual temporal expectation effects were not enhanced by explicit temporal knowledge in a simple visual TE task (Ball et al., 2019). Noteworthy, targets in this experiment were easily perceivable, accuracy was almost at ceiling (only response times were affected) and explicitly provided temporal information was (at least in one explicit group) 100 % valid. Together with reports from other fields, our previous findings suggest that explicit knowledge – when tested in the same experimental context – might not affect performance as compared to implicit statistical learning. However, it has yet to be tested whether the absence of explicit knowledge effects on TE is generalizable across the time course of the experiment (i. e. the learning process), different sensory contexts as well as different, more complex experimental paradigms. Furthermore, we previously argued (Ball, Fuehrmann, et al., 2018; Ball, Michels, et al., 2018) that temporal expectations are not always transferred across individual modalities. However, one aspect of the effects of explicit knowledge might be that it eases the transfer of temporal information between modalities.

In this study, we therefore tested the influence of continuous learning and explicit knowledge about task structure on performance and temporal expectations – within the same paradigm – but under more complex task regimes (multisensory task with distractor stimuli). Here we present a large scale data set (n = 200) based on 4 different complex multisensory experimental designs in which we presented sequences of unisensory (auditory [A] or visual [V]) or multisensory (audiovisual [AV] stimuli with synchronous onsets) stimuli. In Designs 1 and 2 (“easy” task), modality-specific uncertainty was low meaning that with sequence onset, participants knew target’s modality (A sequence with A target, V sequence with V target, AV sequence with AV target). In Designs 3 and 4 (“difficult” task), modality-specific uncertainty was high, i.e. with sequence onset, participants did not know target’s modality (AV sequence with A target, AV sequence with V target, AV sequence with AV target). Target stimuli (A, V, or redundant AV) appeared with a certain probability early or late within the stimulus sequence (11 stimuli presented successively with 1 target embedded among 10 distractors). Likelihood of target occurrence within the sequence (at 3^rd^ or 9^th^ position) was manipulated run-wise, with runs in which early or late targets were more likely.

Based on our previous reports (Ball, Fuehrmann, et al., 2018; Ball, Michels, et al., 2018), which only took into account average performance scores, we hypothesize that performance should be best explained by individual, modality-specific learning curves (modality-specific rule learning). Given that the TE effects were overall more robust in the audiovisual condition, we expected earlier learning trials (i.e. learning curve exceeding chance level) for the audiovisual compared to the unisensory target trials. Additionally, given learning effects – such as in contextual cueing experiments – one might expect that with repeated exposure TE effects might be more pronounced. However, if information is transferred between expected and unexpected runs, TE effects might be rather stable across the experiment, as the TE effect solely depends on re-learning the new target position in each run. Based on our previous publication and participants’ reports, we expected that explicit knowledge might be ineffective to modulate performance and the strength of TE. However, if explicit knowledge eases the transfer of information across modalities, we should find that the learning model best describing the data differs between participants with explicit and implicit temporal knowledge (one learning curve for all modalities vs. modality specific learning curves). Furthermore, explicit knowledge might be associated with earlier learning onsets. Finally, if explicit knowledge modulates behaviour, we should find larger TE effects for participants with explicit knowledge, especially under complex experimental manipulations which can make extraction of temporal regularities more difficult.

## 2. Methods

Note that parts of the data set used here (Exp. 1 to 4, n = 120) were used in previous publications (Ball, Fuehrmann, et al., 2018; Ball, Michels, et al., 2018). To enhance the robustness of statistical estimations, we extended (n = 80) the previous data set with data based on the same experimental design but belonging to so far unpublished data sets. Importantly, we addressed novel research questions whether and how explicit knowledge and learning of temporal regularities affect the strength of temporal expectations, and we focused on estimates derived from a learning model.

### 2.1. Participants

In all experiments, participants were tested after giving signed informed consent. Volunteers who reported any neurological or psychiatric disorders or reduced and uncorrected visual acuity were excluded from the study. Here we used the same exclusion criteria as in our previous reports (Ball, Fuehrmann, et al., 2018; Ball, Michels, et al., 2018): Participants were excluded if they expressed a severe response bias (one response option used in more than 65 % of all trials) and/or performance was well below chance level in one or more conditions (accuracy below 25 %). This study was approved by the local ethics committee of the Otto-von-Guericke-University Magdeburg. In all experiments, we used an independent sample of naïve participants, except for Experiments 6_1 and 6_2 (experiment with 2 sessions per participant) for which samples were almost identical (for demographical data see Table 1). Note that samples were only almost identical as one participant might have been excluded from Experiment 6_1 but not 6_2 based on our exclusion criteria.

**Table 1:** Demographical data of each of the experiments. See Fig.1 for a description of all 4 experimental designs.

| **Experiment** | **Design** | **Mean Age ± SD** | **# Women** | **# left-handed** |
| --- | --- | --- | --- | --- |
| **Exp. 1** | 1 | 24.5 ± 2.7 | 13 | 2 |
| **Exp. 2** | 2 | 23.1 ± 3.4 | 18 | 0 |
| **Exp. 3** | 3 | 24.3 ± 3.6 | 21 | 4 |
| **Exp. 4** | 4 | 23.9 ± 3.7 | 22 | 2 |
| **Exp. 5** | 2 | 23.6 ± 2.7 | 15 | 3 |
| **Exp. 6_1** | 2 | 23.3 ± 4.7 | 22 | 1 |
| **Exp. 6_2** | 4 | 22.9 ± 4.3 | 22 | 1 |

### 2.2. Apparatus

The experiments were programmed using the Psychophysics Toolbox (Brainard, 1997) and Matlab 2012b (Mathworks Inc.). Stimuli were presented on a LCD screen (22’’ 120 Hz, SAMSUNG 2233RZ) with optimal timing and luminance accuracy for vision researches (Wang & Nikolić, 2011). Resolution was set to 1650×1080 pixels and the refresh rate to 60 Hz. Participants were seated in front of the monitor at a distance of 102 cm (eyes to fixation point). Responses were collected with a wireless mouse (Logitech M325).

### 2.3. Stimuli

Uni- or multisensory stimulus sequences (pure tones, circles filled with chequerboards, or a combination of both) were presented on each trial. Chequerboards subtended 3.07° visual angle, and were presented above the fixation cross (centre to centre distance of 2.31°). Sounds were presented from one speaker placed on top of the screen (Designs 1 and 3) at a distance of 7.06° from fixation, 4.76° from chequerboard’s centre, and 3.22° from chequerboard’s edge or via headphones (Sennheiser HD 650; Designs 2 and 4). The speaker was vertically aligned with the centre of the chequerboard stimulus. Chequerboards were presented on a dark grey background (RGB: 25.5). The fixation cross (white) was presented 2.9° above the screen’s centre.

Chequerboards and pure sounds were used as targets and distractors. The distractor frequencies were jittered randomly between 4.6, 4.9, and 5.2 cycles per degree for chequerboards and between 2975, 3000, and 3025 Hz for sounds. Visual and auditory target frequencies were individually adjusted to a 75 % accuracy level at the beginning of the experiment. Hence, targets – although the same type of stimulus (chequerboard/pure sound) – were either lower or higher in frequency compared to distractor frequencies. Furthermore, the intensities for both target and distractor chequerboards and sounds were varied randomly throughout the stimulus sequences. The non-white checkers were jittered between 63.75, 76.5, and 89.25 RGB (average grey value of 76.5 RGB). The sound intensities were jittered between 20 %, 25 %, and 30 % of the maximum sound intensity (average of 25 % = 52 dB[A]). The sound intensity in the experiments with headphones was adjusted to match the sound intensity used for speaker experiments.

### 2.4. Procedure

Participants were seated in a dark, sound-attenuated chamber. For each trial, a sequence consisting of 11 stimuli was presented (see Figure 1 for example sequences). Stimulus duration was 100 ms and stimuli were separated by a 100 ms gap. All stimuli within a sequence were either auditory, visual, or combined auditory and visual stimuli (synchronous presentation). Target stimulus pairs on multisensory trials were always redundant, i.e. targets of both modalities had congruently either a lower or higher frequency than distractors. For each trial, we presented one target stimulus or target stimulus pair (audiovisual sequences) at the 3rd (onset at 400 ms, early target) or 9th position (onset at 1600 ms, late target) of the sequence. Participants were instructed to maintain fixation throughout the experiment and were told that a target was present in each trial. They were required to discriminate the frequency (low or high) of the target as quickly and accurately as possible using a 2-alternative forced-choice procedure. Participants held the mouse with both hands, while placing their left/right thumbs on the left/right mouse buttons, respectively. Each button was used for one of the two response options (key bindings were counterbalanced across participants). The response recording started with the onset of the first stimulus of the sequence and ended 1500 ms after sequence’s offset. Only the first button press was recorded. The end of the response period was then followed by a 200 - 400 ms inter-trial-interval. In case no button was pressed, the trial was repeated (mean of repeated trials across participants: 0.7 % ± 1.4 % SD).

**Figure 1:**
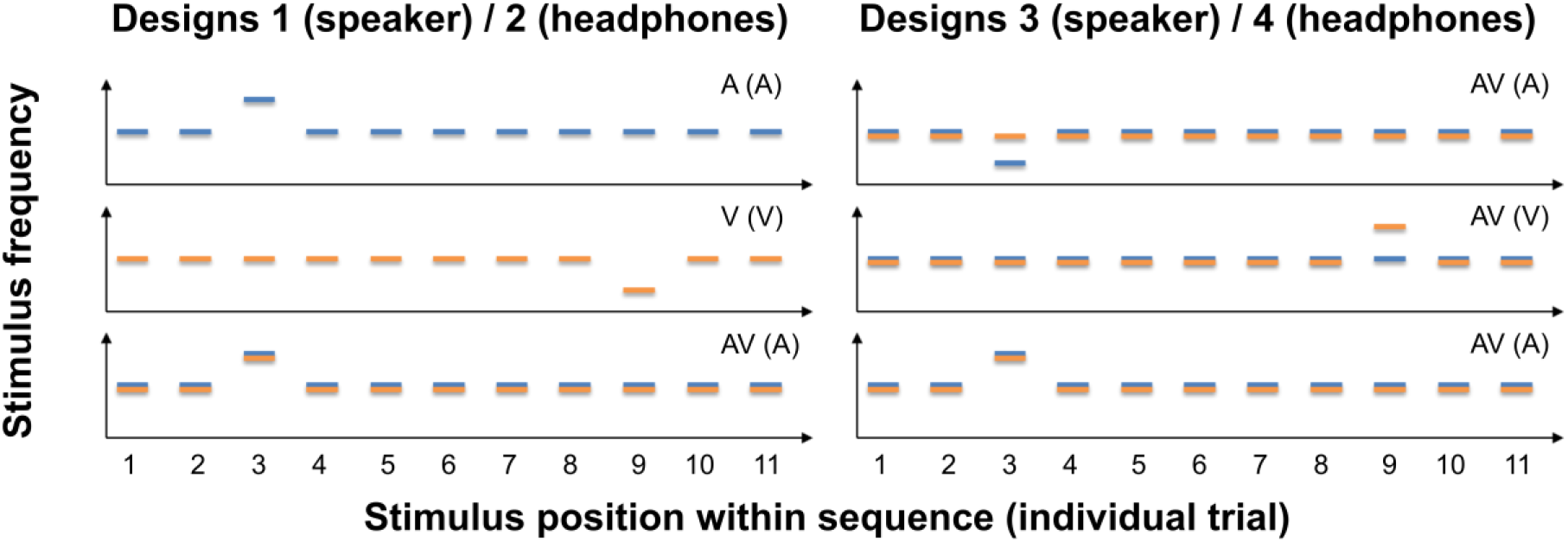
Schematic examples for stimulus sequences in Experimental Designs 1 to 4 (blue = auditory [A], orange = visual [V], blue + orange = audiovisual [AV]). (Left) Stimulus sequences used in Experimental Designs 1 and 2. Each bar represents one of 11 stimuli with a specific stimulus frequency. Bars above or below distractors (auditory distractor frequency: 2975 - 3025 Hz, visual distractor frequency: 4.6 - 5.2 cycles per degree) depict high or low frequency targets, respectively. Targets were only presented at position 3 or 9. The modality of the sequence are depicted in the upper right corner of each graph, with the modality of the target in brackets (e.g. AV (AV) = audiovisual sequence with audiovisual target). **(Right)** In Experimental Designs 3 and 4, we only presented audiovisual sequences, but targets could be either unimodal (A or V) or bimodal (AV). Whenever a target was bimodal, both auditory and visual stimuli were *always* either higher or lower in frequency compared to distractors (redundant target). Note that we only show examples but not all combinations of factors early/late target, higher/lower frequency target and target modality (A, V, AV) which were completely counterbalanced in all experiments.

Each experiment contained three sessions: an initial training session to familiarise participants with the task, a threshold determination session, and the main experimental session. During *‘training’* (24 trials) and *‘threshold determination’* runs (144 trials), we presented unisensory sequences only (auditory or visual). Low and high frequency, early and late occurring, and auditory and visual targets were balanced in these runs. Prior to the experimental sessions, we always conducted two threshold determination runs. After threshold acquisition, visual and auditory stimuli were individually adjusted to 75 % accuracy for all the aforementioned conditions. Each ‘*main experimental session’* was separated into 6 runs (168 trials per run, i.e. 1008 trials total), and we presented all stimulus types (unisensory auditory and visual and multisensory stimuli) and modulated temporal expectation by presenting different numbers of early and late targets within runs. An 86 % likelihood of early target occurrence (always at the 3rd position) and a 14 % likelihood of late targets (9th position) within the stimulus sequence was used for “expect early” runs. In “expect late’’ runs, early target occurrence was reduced to 43 %. We chose this procedure instead of a complete reversal of probabilities in order to obtain a robust estimate of the performance in unexpected early trials as early trials were at the focus of our analysis (see below). Expected and unexpected runs (3 runs each) alternated throughout the experiment, and the type of the first run was counterbalanced across participants. Importantly, all participants were naïve with regard to the changing likelihoods of target position across runs at the beginning of the experiments. Within each run, the number of trials was balanced for sequence types, target types, and target frequencies. Additionally, the number of auditory, visual, and multisensory stimuli, early and late, and low and high targets was balanced across each quarter of runs.

The overall procedure for each experiment (1 to 6_2; for individual design see Table 1) was identical with the following exception: Modality-specific uncertainty was modulated by changing the type of stimulus sequences across experiments. In one experimental design (Designs 1 & 2), we presented either unisensory sequences with unisensory targets or multisensory sequences with multisensory redundant targets (low modality-specific uncertainty). During high modality-specific uncertainty (Designs 3 & 4), we presented only audiovisual sequences, BUT targets were, as before, either unisensory (auditory or visual) with a concurrent distractor in the second modality or redundant multisensory targets (high modality-specific uncertainty). Thus, to perform the task, participants were forced to equally monitor both modalities on each trial to be able to detect the target. Furthermore, auditory stimuli were either delivered over speakers (Designs 1 & 3) or headphones (Designs 2 & 4).

#### 2.4.1. Post-experimental assessment of explicit knowledge

To assess whether participants gained explicit knowledge about temporal structures, we conducted a post-experimental interview. Note that the classification of explicit and implicit knowledge based on verbal reports is a common procedure and has successfully been used in previous studies (Ball et al., 2019; Ewolds, Bröker, de Oliveira, Raab, & Künzell, 2017; Heuer & Schmidtke, 1996; Nissen & Bullemer, 1987). We asked the following questions to gain exhaustive insight into participants’ explicit state of knowledge about temporal regularities. The order of questions was kept constant. Note that early questions were open to avoid biasing, but later questions were more specific to encourage participants to report even vague explicit insights:

- Did you notice any regularities throughout the experiment?
- Was there something specific about the position of target appearance within the sequence?
- Could you please guess at which positions the target was presented.
- Was there a difference between odd and even runs?
- Was there a position pattern throughout the run or was it random across trials?
- Could you please guess the difference between runs.

If participants were unsure about the meaning of ‘trial’ and ‘run’, we further elaborated on the meaning of these terms (trial = one sequence of 11 stimuli for which the frequency of a target had to be reported, run = all trials between two breaks). Only if the answers to all questions were negative, i.e. participants reported the target to appear at all positions or at clearly incorrect positions (e.g. “Target was always presented at the 6^th^ position”), participants were labelled as having only implicit knowledge. Whenever they gave a correct or at least partly correct answer (e.g. “Target was mostly presented early”), they were labelled as having explicit knowledge. We opted for including partly correct answers as exact stimulus positions were difficult/impossible to count given the stimulation frequency (Ball, Fuehrmann, et al., 2018; Ball, Michels, et al., 2018) and participants were at least aware that stimuli occurred mostly early in the sequence (i.e. not the middle or end).

### 2.5. Analysis

In our previous report (Ball, Michels, et al., 2018), we used a criterion to remove response outliers and confirmed – by applying multiple criteria – that the removal of outliers did not affect the results. The present analyses required the same number of trials across participants and conditions to calculate the learning curves. To this end, responses faster than 150 ms were *not excluded* but rather labelled ‘incorrect’ as they are unlikely based on processing of perceptual input. To confirm that this labelling does not change our previous results, we re-ran all our prior analysis (Ball, Michels, et al., 2018). The results (see Supplement 1) are virtually identical to the ones in our previous report. Thus, our previously presented effects appear to be rather robust to the choice of outlier treatment.

Furthermore, as in our previous publications and publications by other groups, we test for temporal expectation effects by focussing on early target trials; late targets are always expected and may thus not require temporal attention (see Ball, Fuehrmann, et al., 2018; Ball, Michels, et al., 2018, for a demonstration of the absence of late target TE effects for the present data set; see also Jaramillo & Zador, 2011; Lange & Röder, 2006; Lange, Rösler, & Röder, 2003; Mühlberg, Oriolo, & Soto-Faraco, 2014 for similar a approach).

#### 2.5.1. Learning model selection

One part of our analyses was based on the logistic regression algorithm developed by Brown and colleagues (Smith & Brown, 2003; Smith et al., 2004). The algorithm (used in Matlab 2017b, Mathworks Inc.) takes into account the binary responses on each trial (correct trials: 1, incorrect trials: 0) and fits a learning curve to the data by using a state-space model of learning, in which a Bernoulli probability model describes behavioural task responses and a Gaussian state equation describes the unobservable learning state process (Kakade & Dayan, 2002; Kitagawa & Gersch, 1996; Smith et al., 2004; Wirth et al., 2003). Furthermore, the algorithm is based on an expectation maximization algorithm (Dempster, Laird, & Rubin, 1977) which computes the maximum likelihood estimate of the learning curve (including the respective confidence intervals).

While the individual learning curves can be used to assess e.g. when a participant has learned the temporal rule (i.e. on trial X), the learning model itself can be used to test different assumptions about whether and how information is transferred across modalities (AV, A, V) and runs. To identify which underlying process best describes the data, we tested three different models by computing the learning curves on different portions of the trials (see Table 2). For instance, if participants learn temporal rules (i.e. target appears at position 3) independently for each modality, we should observe that the best model is based on individual learning curves for each modality (here Model 2; see Table 2). The best model was determined for each participant by calculating the Akaike Information Criterion (AIC) for each model, transforming the AIC into AIC weights and finally calculating the evidence-ratio for each model (Wagenmakers & Farrell, 2004). To foreshadow, Model 2 was on average as well as individually the best model across all experimental designs.

**Table 2:** Description of individual learning models and the related assumptions.

| Model | Learning curves | Assumption |
| --- | --- | --- |
| M1 | learning curve for all early target trials | • information generalised across modalities<br>• information transferred across runs |
| M2 | learning curve for each condition (3 curves total) | • information NOT generalised across modalities<br>• information transferred across runs |
| M3 | learning curve for each condition and for each run (18 curves) | • information NOT generalised across modalities<br>• information NOT transferred across runs |

**Table 3:** Subject-specific best-fitting Model (Model 1 to 3) separated by participants’ temporal awareness. Rows: Number of participants are listed separately for each experiment as well as summarized across experiments (‘Total’). Columns: Number of Participants’ are listed based on the best-fitting-model and participants’ awareness of temporal regularities (explicit and implicit groups) plus total explicit and implicit group size (N). Model 1 (single learning curve), Model 2 (modality-specific learning curves), Model 3 (modality-specific and TE specific learning curves)

| Experiment | Implicit group |  |  |  | Explicit group |  |  |  |
| --- | --- | --- | --- | --- | --- | --- | --- | --- |
|  | N | Model 1 | Model 2 | Model 3 | N | Model 1 | Model 2 | Model 3 |
| Exp. 1 | 18 | 4 | 14 | 0 | 12 | 1 | 11 | 0 |
| Exp. 2 | 21 | 2 | 19 | 0 | 9 | 2 | 7 | 0 |
| Exp. 3 | 20 | 1 | 19 | 0 | 10 | 1 | 9 | 0 |
| Exp. 4 | 24 | 5 | 19 | 0 | 6 | 0 | 6 | 0 |
| Exp. 5 | 15 | 3 | 12 | 0 | 6 | 1 | 5 | 0 |
| Exp. 6_1 | 23 | 2 | 21 | 0 | 7 | 1 | 6 | 0 |
| Exp. 6_2 | 21 | 2 | 19 | 0 | 8 | 2 | 6 | 0 |
| *Total* | *142* | *19* | *123* | *0* | *58* | *8* | *50* | *0* |

#### 2.5.2. Data analyses

Three different performance measures were analysed: mean percent correct, mean reaction times and the ‘learning trial’. The learning trial was specified as the first trial for which the lower confidence interval of the learning curve reliably exceeded chance level (here .5 due to the 2AFC) and stayed above threshold till the end of the experiment (Smith et al., 2004). Learning trials were calculated based on the overall best learning model. For accuracies and RT, we calculated the average scores dependent on factors TE (expected, unexpected), modality (AV, A, V), and run (runs 1&2, runs 3&4, runs 5&6) to test for effects of dynamics of learning (change of TE effect across runs) and modality on TE. Further, we added the factors spatial uncertainty (low, high) and knowledge (implicit, explicit). Note that “modality-specific uncertainty” could not function as a direct interaction term due to the partially crossed design. However, our previous reports revealed that only spatial uncertainty significantly influenced the interaction of modality and TE (Ball, Fuehrmann, et al., 2018; Ball, Michels, et al., 2018), while modality-specific uncertainty contributed the least to TE effects.

For analyses, we used Matlab 2017b (Mathworks Inc.), R (v. 4.0.0) and RStudio (v. 1.0.153). For statistical analyses, we used the ‘afex’ package (v. 0.27-2) in R as data was partially crossed (for factor modality-specific uncertainty). To identify which combination of factors affect TE effects most, we used again a model comparison approach. To this end, we defined all possible interaction models of the factors modality, run, spatial uncertainty and knowledge with factor TE as well as a pure intercept model (see Supplement 2 for overview). To account for the partially crossed data structure, each model also modelled random effects intercepts based on factors modality-specific uncertainty and participant. Hence, we used 16 random intercept models for comparison based on the AIC evidence ratio. For learning trials, we formed all interaction models based on factors modality, knowledge and spatial uncertainty (8 models) for comparison. After determining the best model, individual differences between conditions within the winner model were analysed by using F-values (calculated with the KR-method) and post-hoc t-tests (calculated using the ‘emmeans’ package, v. 1.4.7, with asymptotic dF).

Learning curves were analysed in Matlab by means of cluster permutation tests (for introduction see Maris & Oostenveld, 2007) to identify potential differences between modalities as well as knowledge groups. We used a significance threshold of .05 to identify potential clusters in time (consecutive time points with significant result) and a final cluster-threshold of .004 (Bonferroni corrected to account for multiple testing; note that False-Discovery-Rate correction resulted in the same outcome) to determine differences between conditions. Below we report the corrected cluster-threshold p values (p_cluster_). For the comparison of modalities, we used dependent-sample t-tests and for comparisons of knowledge groups, we used independent sample t-tests to determine the maximum sum t-value of each ‘original data’ cluster as well as the ‘random permutation’ clusters which are used to form the null hypothesis. Note that the cluster test procedure required that learning curves differences are tested separately for low and high modality-specific uncertainty. Hence, we also repeated our model selection approach for the abovementioned data scores (now split for modality-specific uncertainty) to compare it to the outcome of the cluster analyses.

## 3. Results

### 3.1. Results of post-experimental questionnaire

In 58 cases out of the 200 collected data sets, participants noticed any position regularity (Exp. 1: 12, Exp. 2: 9, Exp. 3: 10, Exp. 4: 6, Exp. 5: 6, Exp. 6_1: 7, Exp. 6_2: 8). In 15 cases participants stated regularities immediately. All others only reported regularities after follow-up questioning. 41 participants could identify the second or third position as target position while the remaining stated that targets “occurred mostly early” or “mostly early and late”. Out of the 58 participants, 11 made their statements specifically for the auditory but not visual stimulus.

### 3.2. Learning model selection results

The model that best described the single-trial data across all participants (86.5 % of participants) was Model 2 (see **Fehler! Verweisquelle konnte nicht gefunden werden**.): here the data was modelled independently for each modality while expected and unexpected trials were not further split up. Thus, the best model suggests that information about the most likely temporal position is not generalised across modalities, but that information about the temporal position (early targets) is carried over between runs. Note that this result was independent of explicit knowledge, experiments and design types (see **Fehler! Verweisquelle konnte nicht gefunden werden**.). A depiction of the model evidence ratios is displayed in **Fehler! Verweisquelle konnte nicht gefunden werden**..

### 3.3. Average performance scores analyses – whole data set

The model selection, based on the AIC evidence ratio, suggested a clear winner model for each of the three performance measures, which was also independent of the split into the different modality-specific uncertainty conditions (see Figure 3). For both, the accuracy (Acc) and the RT data, the winning model consisted of the interaction of TE * modality. For the learning trial data, the full model (modality * spatial uncertainty * knowledge) accounted best for the data at hand.

**Figure 2:**
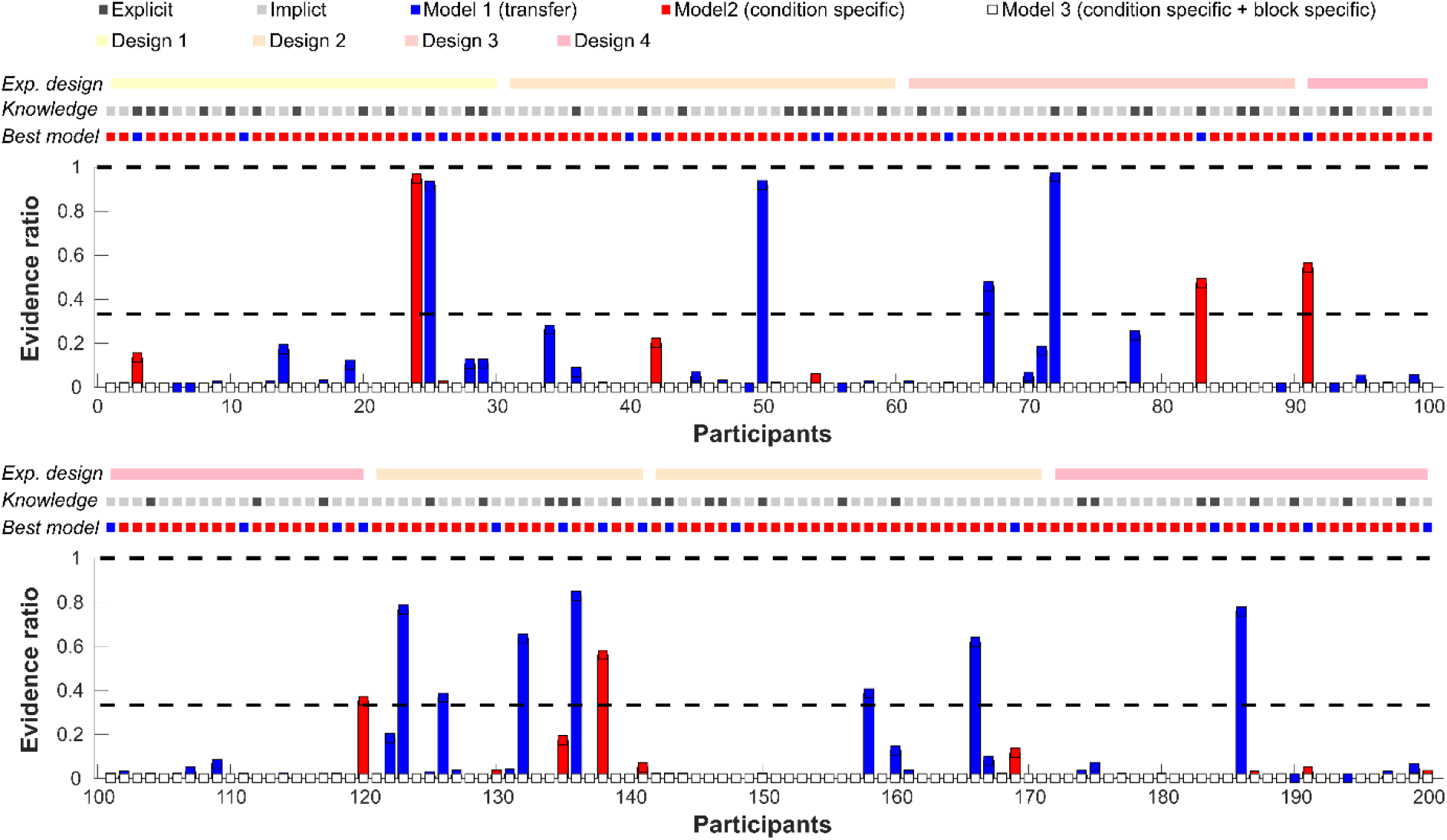
Evidence ratio for each learning model (based on estimated learning curves). Each number on the x-axis represents an individual participant. Top: Participants 1-100 (Exp. 1 to 4). Bottom: participants 101-200 (Exp. 4 to 6_2). The top dotted lines indicate the experimental design (*Design 1*: low spatial and low modality-specific uncertainty, *Design 2*: high spatial and low modality-specific uncertainty, *Design 3*: low spatial and high modality-specific uncertainty, *Design 4*: high spatial and high modality-specific uncertainty), participants’ knowledge (light grey = implicit, dark grey = explicit) and the individual best model (blue = Model 1 (single learning curve), red = model 2 (modality-specific learning curves), white = Model 3 (modality-specific and TE specific learning curves)). Please note that Model 3 was never the best Model. The bar graph below depicts individual evidence ratios for the second and third best models as compared to the best model. A score of .5 e.g. implies that the best model was two times more likely than the other model. Here we highlighted the .33 score as this value indicates that the best model was 3 times more likely than the all other models (in close resemblance to the Bayes factor).

**Figure 3:**
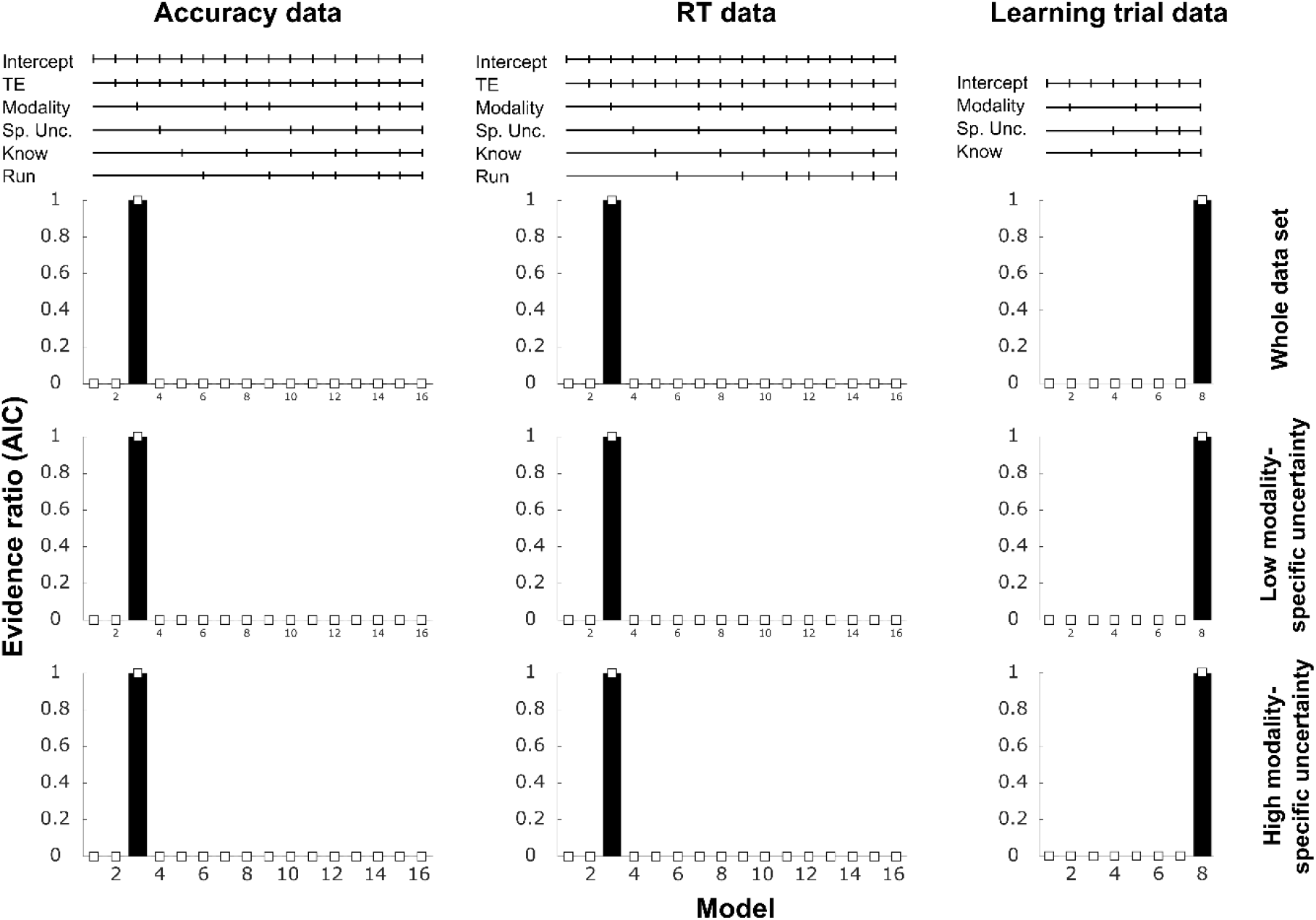
Comparison of mixed-models for each average performance score based on AIC evidence ratio.). Accuracy (left column), RTs (middle column) and learning trial (right column). The top line graphs depict the factors that were included as interaction terms in the respective model (Sp. Unc. = spatial uncertainty, Know = knowledge group). Bar graphs below show the evidence ratios for analyses of the whole data set (top row) and the split into low and high modality-specific uncertainty (middle and bottom row). Note that the best model has an evidence ratio of 1 (best model score divided by itself = 1); and that, for instance, a score of .333 implies that the comparison model was three times less likely than the best model. More importantly, the graph shows that all ‘non-best’ models showed evidence ratios close to zero (as indicated by the white square markers close to the x-axis), indicating that these models are highly unlikely as compared to the best model.

Following up on the individual differences in estimated marginal means, we found a significant main effect of TE (Acc: F(1, 3442.16) = 70.21, p < .001; RT: F(1, 3442.16) = 274.62, p < .001), a main effect of modality (Acc: F(2, 3442.16) = 241.56, p < .001; RT: F(2, 3442.16) = 264.63, p < .001) as well as an interaction of both factors (Acc: F(2, 3442.16) = 3.8, p = .022; RT: F(2, 3442.16) = 3.0, p = .05) for both the accuracy and RT data. Summarized, participants responded more often correctly and faster in the expected compared to the unexpected condition and this TE effect was enhanced in the AV and A conditions as compared to the V condition (see Figure 4, top two rows). For the learning trials, only the factor modality was significant (F(2,422.99) = 16.41, p < .001) with earlier learning trials in the AV compared to the A and V conditions (see Figure 4, 3^rd^ row). All other effects were non-significant (F < 2.5, p > .115).

**Figure 4:**
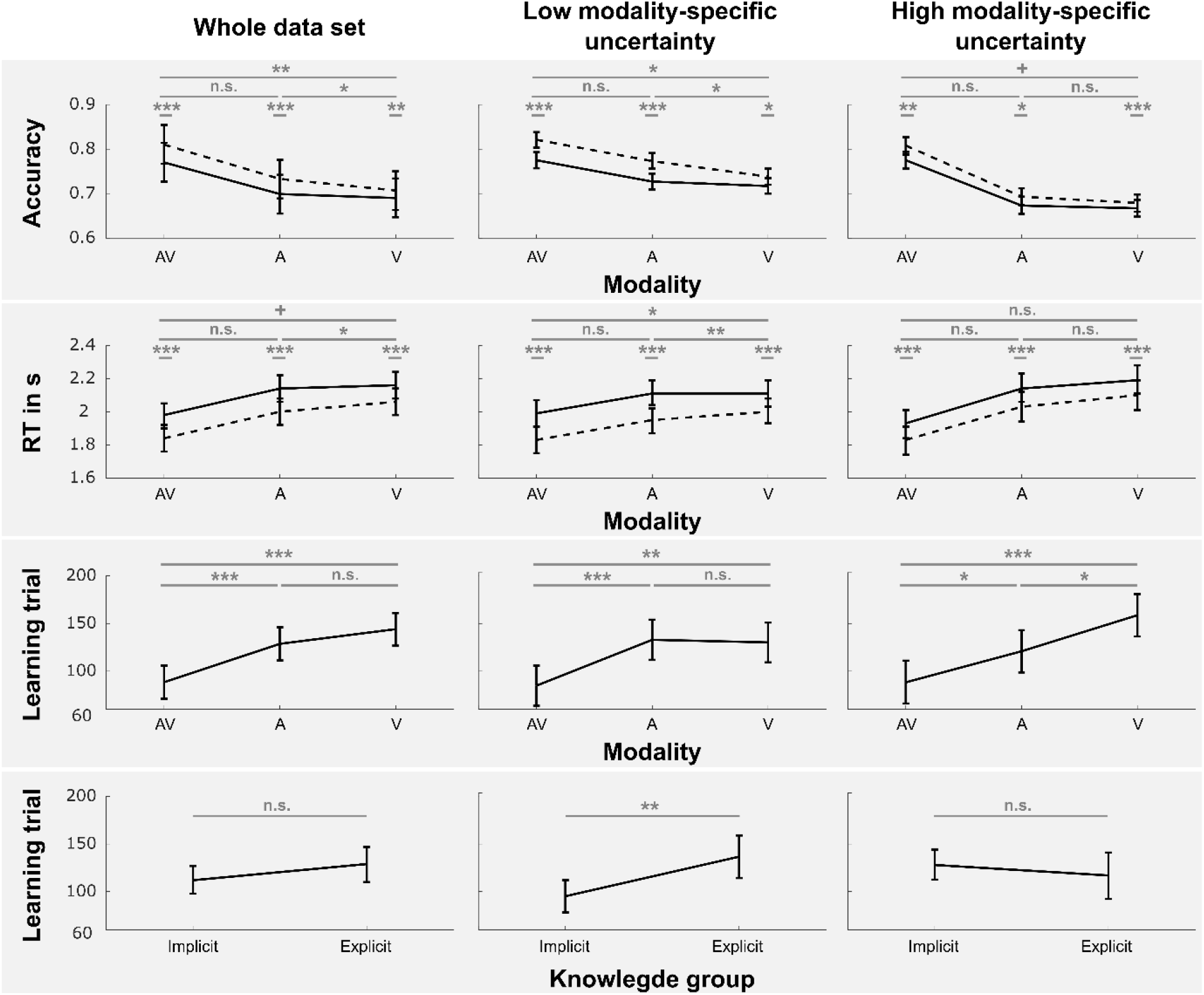
Group-mean scores (estimated marginal means) for accuracy (1^st^ row), RT (2^nd^ row) and learning trial (3^rd^ – 4^th^ row) effects. Averages are presented for analyses of the whole data set (left) as well as for the split into low (middle) and high modality-specific uncertainty (right). Statistical comparisons (uncorrected) are labelled as follows: n.s. (p > .1), + (p>.05), * (p>.01), ** (p>.001), *** (p>=0). Note that on the top two rows, data is split into expected (dotted line) and unexpected (solid line) trials. Further, statistical results are based on testing the difference between expected and unexpected trials as well as testing whether this difference was altered by the modality context. Row 3-4 displays modality- and knowledge-specific effects of the learning trial.

### 3.4. Average performance scores analyses – Low modality-specific uncertainty (known target modality)

The model selections for accuracy and RT data, based on low modality-specific uncertainty only, were in line with the model selection for the whole data set (see Figure 4, middle). The best model in both cases was model 2 (see Figure 3, 2^nd^ row), which was based on the interaction of TE and modality. Again, we found significant effects of TE (Acc: F(1,1882) = 64.06, p < .001; RT: F(1,1882) = 229.97, p < .001), modality (Acc: F(2,1882) = 77.67, p < .001; RT: F(2,1882) = 90.74, p < .001) as well as the interaction of both terms (Acc: F(2,1882) = 3.23, p = .04; RT: F(2,1882) = 4.34, p = .013) with essentially the same results pattern as for the whole data set. For the learning trials, the results were also virtually identical (modality: F(2,214) = 7.64, p < .001; see Figure 4, 3^rd^ row). However, there was also a significant effect of knowledge (F(1,107) = 8.4, p = .005; all other effects F < 2.38, p > .095), with earlier learning trials for the implicit compared to the explicit group (see Figure 4, 4^th^ row).

The analyses of learning curves mirrored the results for the accuracy data (note that we did not split data for low and high spatial uncertainty, based on the outcome of the mixed model comparison procedure). First of all, there was a clear cut modality effect: performance was increased in the AV compared to the A (1 cluster with p_Cluster_ = 0) and V conditions (1 cluster with p_Cluster_ = 0). Although, cluster time to be interpreted cautiously (Sassenhagen & Draschkow, 2019), the cluster results suggest that ranges have conditions differed across the whole time range of the experiment. Thus, differences between conditions were driven independent of learning. No differences were found between the A and V conditions (all p_Cluster_ > .252). For the comparison of participants with explicit and implicit knowledge, no significant differences were found (all p_Cluster_ > .468). More importantly, even if the groups would differ, performance would be lower in the explicit knowledge group – the opposite of the effect one would expect for the influence of explicit knowledge on behaviour (see Figure 5). Additionally, the results in Figure 5 strongly suggest that after the initial onset, performance varied only minimally across time.

**Figure 5:**
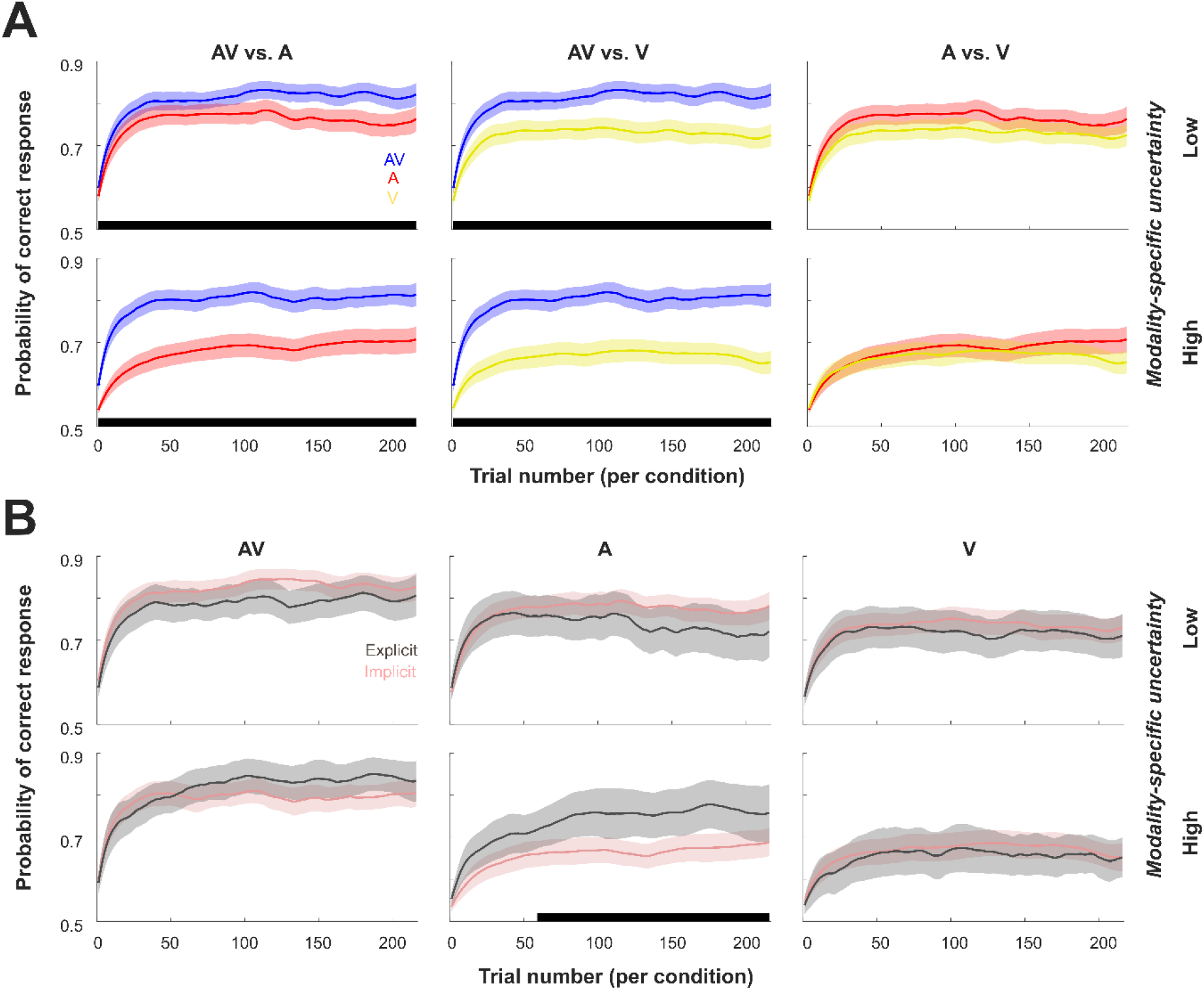
Learning curve data comparison for modalities (A) and knowledge groups (B). Time ranges of significant clusters (consecutive significant differences across trials between conditions) are highlighted with black bars. Results are presented for low (top rows) and high (bottom rows) modality-specific uncertainty. Colours used for conditions: AV – blue, A – red, V – yellow, Explicit group – dark grey, Implicit group – light red.

### 3.5. Average performance scores analyses – High modality-specific uncertainty (unknown target modality)

Again, Model 2 was the best-fitting model for the accuracy and RT data and Model 8 was the best-fitting model for the learning trials (see Figure 3, 3^rd^ row). Note that there were two differences to the abovementioned results. First, the interactions of TE * modality did not reach significance under high modality-specific uncertainty (Acc: F(2,1508) = 1.39, p = .25; RT: F(2,1882) = F(2,1508) = .249, p = .799) and second, the learning trials were not only faster between the AV and unisensory conditions but also for the A compared to the V condition (F(2,170) = 11.26, p < .001; see Figure 4, 3^rd^ row). Besides the significant TE and modality effects (all p < .001) for accuracies and TE, all remaining effects related to the learning trials were non-significant (F’s < 1.66, p’s > .193).

Similar to results from the low uncertainty condition, analyses of the learning curves showed again a significant modality effect (see Figure 5), with performance being higher in the AV condition compared to both unisensory conditions (p_Cluster_ = 0). There were again no significant differences between the A and V condition (p_Cluster_ > .756). For the comparison of participants with explicit and implicit knowledge, there was a significant difference in the auditory condition (p_Cluster_ = .046). The cluster was spanning the mid to end trials of the experiment, with higher performance in the explicit group. All other differences were non-significant (all p_Cluster_ > .78).

## 4. Discussion

Rule learning - including rules based on temporal probabilities (most likely time point of target occurrence) - and explicit knowledge about temporal rules (e.g. due to temporal cueing) are often assumed to maximise performance by actively preparing for certain events in time (Nobre & Rohenkohl, 2014). However, previous studies typically utilized unisensory-specific contexts. In this study we tested, whether *explicit knowledge* of temporal regularities would affect temporal expectancies (TE) in different uni- and multisensory contexts. Furthermore, we tested whether dynamic learning models can be successfully applied to analyse temporal rule learning and the temporal dynamics of TE effects.

We found that temporal expectations – in the present study – were most likely to be altered by the modality-related context using a direct model comparison approach. Additional factors such as the dynamics of learning (change of TE effect across runs) and explicit knowledge appear to contribute only minimally to the size of TE effects when considering accuracy and reaction times. In accord, these findings were further corroborated by parameters derived from learning models. In particular, the probability of answering correctly was clearly different across modalities but this difference was stable across all trials (i.e. no modality-specific differences due to learning occurred over time). Further, no consistent overall effect was found for the differences between knowledge groups. Explicit knowledge only improved performance for auditory condition under high modality-specific uncertainty but was also linked to slower learning (shallower learning curves) under low modality-specific uncertainty. Finally, the learning model suggested that performance was modality-dependent; thus, most participants showed no rule transfer across conditions.

State-based learning models have previously been used to successfully determine dynamic learning in contexts such as memory-association tasks, visuo-motor associative learning tasks and location-scene association (Clarke et al., 2018; Hargreaves et al., 2012; Smith et al., 2004; Wirth et al., 2003). In this study, we evaluated if they can successfully be applied to temporal rule learning tasks. Our results indicate that differences between the modelled learning curves, closely resemble effects based on measured average performance. Further, we used the model to formally test our assumption that information about the most likely temporal position is not transferred between modalities (Ball, Michels, et al., 2018). The model supported this claim, even on the individual level (86.5 % of all participants) and further suggested that explicit knowledge does not ease the transfer of temporal position knowledge (i.e. the most likely temporal position) across modalities. Importantly, the learning curves also indicate that participants learned the temporal rule faster in the multisensory compared to unisensory contexts thereby extending the findings of previous reports showing increased performance and TE effects in the multisensory condition (Ball, Fuehrmann, et al., 2018; Ball, Michels, et al., 2018). Potentially, the higher informational content of multisensory stimulation – which has been linked to multisensory performance benefits (e.g. Alais & Burr, 2004; Ball, Fuehrmann, et al., 2018; Ball, Michels, et al., 2018; Driver & Noesselt, 2008; Noesselt et al., 2007, 2010; Parise et al., 2012; Starke et al., 2017; Werner & Noppeney, 2010) – also increases speed of extraction of temporal regularities and temporal perceptual learning. Together, the results suggest that learning models can provide additional insights into temporal learning, unravel potentially different learning types (if present; here modality-specific learning) and might also be used to identify the neural origins of temporal rule learning on a trial-by-trial basis in future neuroimaging studies, as has previously been done in other non-temporal research fields (Clarke et al., 2018; Hargreaves et al., 2012; Smith et al., 2004; Wirth et al., 2003).

Turning to the differences between explicit and implicit temporal knowledge, previous research suggests that temporal expectations are created implicitly from the temporal trial structure (e.g. rhythms) and from the overall statistical distribution (e.g. increased likelihood for events with a particular foreperiod in one run) of likely events in time by means of statistical learning (Ball, Fuehrmann, et al., 2018; Ball et al., 2019; Ball, Michels, et al., 2018; Breska & Deouell, 2014; Cravo, Rohenkohl, Wyart, & Nobre, 2013; Rohenkohl et al., 2011; Shen & Alain, 2012). Our results support this assumption as temporal expectation effects in our study were largely driven by implicit knowledge (142 out of 200 participants). Thus, statistics about predictable events – at least in our study – seem to be automatically extracted in the majority of participants and to be utilized independently of explicit knowledge about the temporal manipulation.

To our best knowledge, not many studies have directly compared the effects of explicit vs. implicit knowledge within the same paradigm. However, a difference in behaviour or neural activation patterns – when based on different experimental paradigms or studies without assessment of participants’ explicit knowledge – would be insufficient to conclude that explicit knowledge truly affects e. g. performance and learning patterns (Ball et al., 2019); and most within-study comparisons of explicitly- vs. implicitly-driven behaviour report the absence of modulating effects due to explicit knowledge. For instance, Max and colleagues (2015) investigated the impact of distracting sounds (deviant pitch) within a sequence of standard sounds on auditory duration judgments. The authors found no evidence that instructions (explicit vs. no-information) altered behavioural performance related to the distraction effect, though they observed changes in the EEG signals (i.e. a lower P3a amplitude when distractors were expected due to instructions). They suggest that differences in neural activity index participants’ involuntary shift of attention to distracting sounds which is altered by prior knowledge (implicit vs. explicit instruction) yet these neural activity changes do not necessarily change the behavioural outcome. In addition, the absence of influences of explicit knowledge on performance was also reported in priming studies (Francken, Gaal, & de Lange, 2011; Van den Bussche et al., 2013), contextual cueing studies (Chun & Jiang, 2003; Geyer, Baumgartner, Müller, & Pollmann, 2012; Preston & Gabrieli, 2008; Westerberg, Miller, Reber, Cohen, & Paller, 2011) and motor sequence learning tasks (Sanchez & Reber, 2013). Remarkably, results from some studies even indicate that explicit compared to implicit knowledge (1) can be detrimental (Green & Flowers, 2003; Preston & Gabrieli, 2008; Van den Bussche et al., 2013), (2) reduces differences between e.g. cued and uncued trials (Schlagbauer, Muller, Zehetleitner, & Geyer, 2012), (3) might only be beneficial on a long-term but not short-term basis (Ewolds et al., 2017) or under very specific task constraints (Stefaniak, Willems, Adam, & Meulemans, 2008), and (4) could in principle reflect changes of response strategy rather than a perceptual facilitation (Schlagbauer et al., 2012; Summerfield & Egner, 2009). Recently, we found support for the latter idea of response strategy shifts in a simple visual temporal expectation paradigm (Ball et al., 2019). Here we extent our previous findings to multisensory temporal expectations paradigms by showing that explicit knowledge has only a marginal effect on performance, which is also not always favourable.

In general, we only found weak evidence for differences between participants with explicit and implicit knowledge. In the experiments with high modality-specific uncertainty, accuracies were enhanced in the explicit group, but only in the auditory condition as determined by the learning curve analysis. Note that our previous reports showed that the unimodal auditory condition, was the preferred condition (i.e. unimodal condition with best performance across all unimodal conditions) for most of participants (Ball, Fuehrmann, et al., 2018). Thus, explicit knowledge might potentially help to overcome response conflicts by identifying and discriminating the unisensory target in a multisensory stimulus sequence, but only for the preferred unisensory modality. In accord, 19 % of the explicit participants made their statement specifically about the auditory condition lending further support to the notion that learning occurred modality-specific. Given our results, it is possible that explicit knowledge is ineffective whenever there is no conflict (audiovisual target) or the unisensory temporal position is less known or even unknown (visual target). Another possibility is that explicit knowledge enhances processing of the stimulus, but only in sensory systems with high temporal acuity (i.e. the auditory system; see e.g. Ball, Michels, et al., 2018; Nobre & Rohenkohl, 2014 for discussion of thie issue). However, if this would be true, we would expect to find similar results for the auditory targets under low modality-specific uncertainty, which was not the case. In contrast, the learning curves for the low modality-specific uncertainty experiments even showed a trend for lower performance in the explicit knowledge group. Following up on this pattern of results, it might be that explicit knowledge does mainly help to *detect* the target more reliably in time – thereby increasing the confidence about target presence at certain points in time – while explicit knowledge does (on average) not help to finally *discriminate* the target more reliably. However, such increase in confidence might not always be beneficial and result in premature responses, thereby sacrificing accuracy in discrimination tasks, especially under easy task regimes; an interpretation which would be in line with our previous report (Ball et al., 2019) and other findings (Schlagbauer et al., 2012; Summerfield & Egner, 2009). Under more difficult task regimes, explicit knowledge might help to better cope with the task, at least for specific unisensory conditions and independently of the run-wise manipulation of temporal regularities. Optimally, future tasks would incorporate a measure of detection as well as discrimination performance (without priming the temporal task component) and potentially also confidence ratings to further elucidate the interplay of confidence, detection, discrimination and explicit knowledge in TE tasks. Additionally, the differences between groups appear to be context sensitive – they depend on the modality-context as well as task-difficulty or rather task design. Thus, future studies should also systematically study the influence of these factors to allow forming theories about how TE, modality, distraction and explicit knowledge holistically affect performance.

A possible limitation of the current study is the assessment of implicit and explicit knowledge of temporal regularities. In other research fields like contextual cueing (Chun & Jiang, 1998), explicit knowledge is typically tested by comparing recognition accuracy of old vs. new scenes. However, it is currently debated whether these post-hoc detection tests truly supports the null hypothesis (i.e. implicit knowledge) or whether the assessments of implicit vs. explicit knowledge by means of recognition tests in previous studies were underpowered (Vadillo, Konstantinidis, & Shanks, 2016). Furthermore, it is difficult to design a perceptual post-hoc test for recognizing certain old vs. new temporal target positions as individual temporal positions (e.g. 3rd vs. 5^th^ position in stimulus stream) are hardly distinguishable due to the fast presentation frequency (Ball, Michels, et al., 2018). Instead, we opted for using an exhaustive post-hoc questionnaire (Ball et al., 2019; Ewolds et al., 2017; Heuer & Schmidtke, 1996; Nissen & Bullemer, 1987) and motivated participants to even guess the most likely target positions. Nonetheless, this procedure still does not allow to pin-point the exact time when participants gathered explicit knowledge. To quantify the onset of learning, we chose a computational model which was specifically designed to have higher sensitivity in capturing the ‘true’ learning onset and to test for difference across conditions (Smith et al., 2004); further, the model was successfully applied in previous non-temporal studies (Clarke et al., 2018; Hargreaves et al., 2012; Smith et al., 2004; Wirth et al., 2003). However, assuming that the learning trial represents the speed of learning, we did not find evidence for earlier learning onsets in the explicit group. Thus, if the learning onset in the explicit group represents participants becoming aware of the temporal position, this did not happen faster than implicit learning of temporal position in the implicit group. Hence, this finding might imply that learning onsets in both groups were rather based on implicit statistical learning and that participants noticed the temporal regularity later on in the experiment.

We hypothesized that temporal rule learning might elevate performance with repeated exposure to temporal regularities (change of the size of TE across runs); this assumption was based on results from contextual cueing, perceptual and motor sequence learning studies (Bueti & Buonomano, 2014; Chun & Jiang, 1998; Clegg, Digirolamo, & Keele, 1998). They reported improved performance for repeated (i.e. expected) compared to new (i.e. unexpected) layouts and sequences. However, TE effects were not sufficiently modulated by repeated exposure, neither for response times nor accuracies, as indexed by the model comparison. This might be due to an information transfer across runs. Hence, when participants learned to attend the early position in one run, they slowly shifted their attention away from the early position in the next run, resulting in similar TE effect sizes across runs. Moreover, learning curves only showed minimal accuracy fluctuations over time (see learning curve results). Most importantly, the largest effect we found – the performance difference between modalities – was present in each trial; hence, this difference between modalities did not require to learn e.g. a certain target position. We further hypothesised that explicit knowledge might be based on faster learning, however, as mentioned above, learning speed was similar in the explicit and implicit group. Thus, it appears that explicit rule learning as well as TE effects (based on shifts of attention due to run-wise probability manipulations) are not affected by run-wise dynamics of learning over the course of an experiment. At this point it remains open whether learning effects on TEs are e.g. more pronounced in long-term studies (experiments over several weeks). Nevertheless, based on our data we conclude that short-term adaptation has little effect on temporal expectations in complex task designs.

## 5. Conclusion

Here we show, using a computational learning model, that temporal rules were learned separately for each modality and that explicit knowledge did not ease the transfer of information transfer across modalities. Thus, knowing when a target is likely to occur is not automatically generalised. Further, it is often suggested that explicit knowledge changes the recruitment of cognitive resources. However, our results suggest that top-down modulations of behaviour due to explicit knowledge appear to be detrimental under low task difficulty (in line with previous reports) and only facilitated unisensory auditory target performance when the target modality in a given trial was unpredictable (high task difficulty). Thus, explicit knowledge might partially help to resolve response conflicts under high uncertainty but might render participants overconfident (while sacrificing accuracy) when targets are more easily detectable. Additionally, audiovisual stimulation resulted in faster and better learning. However, dynamics of temporal learning (the size of temporal expectation across runs) only contributed little to the observed TE effects. Together, the results suggest that temporal rule learning might be more reliable in multi-than unisensory context and that explicit knowledge as well as dynamics of learning have little to no beneficial effect on temporal expectation. Given our sample size and the comparatively small effects, our results also stress the importance to carefully interpret changes in behaviour due to explicit knowledge when assessed by across-study reviews.

## Abbreviation

TE: Temporal Expectation
AV/A/V: Audio-Visual / Auditory / Visual
Acc: Mean Accuracy
RT: Reaction Time

## Acknowledgements

We thank L. E. Michels, F. Fuehrmann, F. Stratil, and J. Meiners for acquiring the data. We thank P. Vavra for his helpful comments on our analyses.

## Declarations

### CRediT author statement

**Felix Ball:** Conceptualization, Methodology, Formal analysis, Investigation, Writing - Original Draft, Writing - Review & Editing, Visualization, Supervision, Project administration, Funding acquisition **Inga Spuerck:** Writing - Original Draft, Writing - Review & Editing **Toemme Noesselt:** Conceptualization, Methodology, Writing - Original Draft, Writing - Review & Editing, Supervision, Funding acquisition

### Ethics approval

This study was approved by the local ethics committee of the Otto-von-Guericke-University, Magdeburg.

### Consent to participate and publish

Informed consent was obtained from all individual participants included in the study.

### Open Practices Statement

The data and materials are available in the supplementary information files and upon request (e.g. codes). None of the experiments were preregistered.

### Data availability statement

The authors declare that the analysed data as well as potential supplementary analyses (not reported in the main manuscript) are available in the supplementary information files.

### Code availability

Data were analysed with R/Rstudio (freely available) and the Learning Curve Analysis software package for Matlab (http://www.neurostat.mit.edu/behaviorallearning). The remaining code is available upon request.

## References

Alais, D., & Burr, D. (2004). The Ventriloquist Effect Results from Near-Optimal Bimodal Integration. Current Biology, 14(3), 257–262. https://doi.org/10.1016/j.cub.2004.01.029

Ball, F., Fuehrmann, F., Stratil, F., & Noesselt, T. (2018). Phasic and sustained interactions of multisensory interplay and temporal expectation. Nature Scientific Reports, 8, 10208. https://doi.org/10.1038/s41598-018-28495-7

Ball, F., Groth, R.-M., Agostino, C. S., Porcu, E., & Noesselt, T. (2019). Explicitly vs. implicitly driven temporal expectations: No evidence for altered perceptual processing due to top-down modulations. Attention Perception & Psychophysics, In press.

Ball, F., Michels, L. E., Thiele, C., & Noesselt, T. (2018). The role of multisensory interplay in enabling temporal expectations. Cognition, 170, 130–146. https://doi.org/10.1016/j.cognition.2017.09.015

Batterink, L. J., Reber, P. J., Neville, H. J., & Paller, K. A. (2015). Implicit and explicit contributions to statistical learning. Journal of Memory and Language, 83, 62–78. https://doi.org/10.1016/j.jml.2015.04.004

Brainard, D. H. (1997). The Psychophysics Toolbox. Spatial Vision, 10(4), 433–436. https://doi.org/10.1163/156856897X00357

Breska, A., & Deouell, L. Y. (2014). Automatic bias of temporal expectations following temporally regular input independently of high-level temporal expectation. Journal of Cognitive Neuroscience, 26(7), 1555–1571. https://doi.org/10.1162/jocn_a_00564

Bueti, D., & Buonomano, D. V. (2014). Temporal Perceptual Learning. Timing & Time Perception, 2(3), 261–289. https://doi.org/10.1163/22134468-00002023

Capizzi, M., Sanabria, D., & Correa, Á. (2012). Dissociating controlled from automatic processing in temporal preparation. Cognition, 123(2), 293–302. https://doi.org/10.1016/j.cognition.2012.02.005

Chun, M. M., & Jiang, Y. (1998). Contextual Cueing: Implicit Learning and Memory of Visual Context Guides Spatial Attention. Cognitive Psychology, 36(1), 28–71. https://doi.org/10.1006/cogp.1998.0681

Chun, M. M., & Jiang, Y. (2003). Implicit, long-term spatial contextual memory. Journal of Experimental Psychology. Learning, Memory, and Cognition, 29(2), 224–234. Retrieved from http://www.ncbi.nlm.nih.gov/pubmed/12696811

Clarke, A., Roberts, B. M., & Ranganath, C. (2018). Neural oscillations during conditional associative learning. NeuroImage, 174, 485–493. https://doi.org/10.1016/j.neuroimage.2018.03.053

Clegg, B. A., Digirolamo, G. J., & Keele, S. W. (1998). Sequence learning. Trends in Cognitive Sciences, 2(8), 275–281. https://doi.org/10.1016/S1364-6613(98)01202-9

Correa Á., Cona, G., Arbula, S., Vallesi, A., & Bisiacchi, P. (2014). Neural dissociation of automatic and controlled temporal preparation by transcranial magnetic stimulation. Neuropsychologia, 65, 131–136. https://doi.org/10.1016/j.neuropsychologia.2014.10.023

Correa, Á., Lupiáñez, J., & Tudela, P. (2005). Attentional preparation based on temporal expectancy modulates processing at the perceptual level. Psychonomic Bulletin & Review, 12(2), 328–334. Retrieved from http://www.ncbi.nlm.nih.gov/pubmed/16082814

Coull, J. T., Frith, C. D., Büchel, C., & Nobre, A. C. (2000). Orienting attention in time: behavioural and neuroanatomical distinction between exogenous and endogenous shifts. Neuropsychologia, 38(6), 808–819. Retrieved from http://www.ncbi.nlm.nih.gov/pubmed/10689056

Coull, J. T., & Nobre, A. C. (2008, April). Dissociating explicit timing from temporal expectation with fMRI. Current Opinion in Neurobiology. https://doi.org/10.1016/j.conb.2008.07.011

Cravo, A. M., Rohenkohl, G., Wyart, V., & Nobre, A. C. (2013). Temporal Expectation Enhances Contrast Sensitivity by Phase Entrainment of Low-Frequency Oscillations in Visual Cortex. Journal of Neuroscience, 33(9), 4002–4010. https://doi.org/10.1523/JNEUROSCI.4675-12.2013

de la Rosa M. D., Sanabria, D., Capizzi, M., & Correa, Á. (2012). Temporal Preparation Driven by Rhythms is Resistant to Working Memory Interference. Frontiers in Psychology, 3, 308. https://doi.org/10.3389/fpsyg.2012.00308

Dempster, A. P., Laird, N. M., & Rubin, D. B. (1977). Maximum Likelihood from Incomplete Data Via the EM Algorithm. Journal of the Royal Statistical Society: Series B (Methodological), 39(1), 1–22. https://doi.org/10.1111/j.2517-6161.1977.tb01600.x

Driver, J., & Noesselt, T. (2008, January 10). Multisensory Interplay Reveals Crossmodal Influences on “Sensory-Specific” Brain Regions, Neural Responses, and Judgments. Neuron. https://doi.org/10.1016/j.neuron.2007.12.013

Ewolds, H. E., Bröker, L., de Oliveira, R. F., Raab, M., & Künzell, S. (2017). Implicit and Explicit Knowledge Both Improve Dual Task Performance in a Continuous Pursuit Tracking Task. Frontiers in Psychology, 8, 2241. https://doi.org/10.3389/fpsyg.2017.02241

Fairhurst, M. T., Travers, E., Hayward, V., & Deroy, O. (2018). Confidence is higher in touch than in vision in cases of perceptual ambiguity. Scientific Reports, 8(1), 15604. https://doi.org/10.1038/s41598-018-34052-z

Francken, J. C., Gaal S. van, & de Lange, F. P. (2011). Immediate and long-term priming effects are independent of prime awareness. Consciousness and Cognition, 20(4), 1793–1800. https://doi.org/10.1016/j.concog.2011.04.005

Geyer, T., Baumgartner, F., Müller, H. J., & Pollmann, S. (2012). Medial temporal lobe-dependent repetition suppression and enhancement due to implicit vs. explicit processing of individual repeated search displays. Frontiers in Human Neuroscience, 6, 272. https://doi.org/10.3389/fnhum.2012.00272

Giordano, A. M., McElree, B., & Carrasco, M. (2009). On the automaticity and flexibility of covert attention: a speed-accuracy trade-off analysis. Journal of Vision, 9(3), 30.1-10. https://doi.org/10.1167/9.3.30

Green, T. D., & Flowers, J. H. (2003). Comparison of implicit and explicit learning processes in a probabilistic task. Perceptual and Motor Skills, 97(1), 299–314. https://doi.org/10.2466/pms.2003.97.1.299

Hannula, D. E., & Greene, A. J. (2012). The hippocampus reevaluated in unconscious learning and memory: at a tipping point? Frontiers in Human Neuroscience, 6, 80. https://doi.org/10.3389/fnhum.2012.00080

Hargreaves, E. L., Mattfeld, A. T., Stark, C. E. L., & Suzuki, W. A. (2012). Conserved fMRI and LFP signals during new associative learning in the human and macaque monkey medial temporal lobe. Neuron, 74(4), 743–752. https://doi.org/10.1016/j.neuron.2012.03.029

Henke, K. (2010). A model for memory systems based on processing modes rather than consciousness. Nature Reviews. Neuroscience, 11(7), 523–532. https://doi.org/10.1038/nrn2850

Heuer, H., & Schmidtke, V. (1996). Secondary-task effects on sequence learning. Psychological Research, 59(2), 119–133. https://doi.org/10.1007/BF01792433

Jaramillo, S., & Zador, A. M. (2011). The auditory cortex mediates the perceptual effects of acoustic temporal expectation. Nature Neuroscience, 14(2), 246–251. https://doi.org/10.1038/nn.2688

Kakade, S., & Dayan, P. (2002). Acquisition and extinction in autoshaping. Psychological Review, 109(3), 533–544. https://doi.org/10.1037/0033-295x.109.3.533

Kitagawa, G., & Gersch, W. (1996). Smoothness Priors Analysis of Time Series. Springer New York.

Kurtz, P., Shapcott, K. A., Kaiser, J., Schmiedt, J. T., & Schmid, M. C. (2017). The Influence of Endogenous and Exogenous Spatial Attention on Decision Confidence. Scientific Reports, 7(1), 6431. https://doi.org/10.1038/s41598-017-06715-w

Lange, K., & Röder, B. (2006). Orienting attention to points in time improves stimulus processing both within and across modalities. Journal of Cognitive Neuroscience, 18(5), 715–729. https://doi.org/10.1162/jocn.2006.18.5.715

Lange, K., Rösler, F., & Röder, B. (2003). Early processing stages are modulated when auditory stimuli are presented at an attended moment in time: an event-related potential study. Psychophysiology, 40(5), 806–817. Retrieved from http://www.ncbi.nlm.nih.gov/pubmed/14696734

Mancini, F., Dolgevica, K., Steckelmacher, J., Haggard, P., Friston, K., & Iannetti, G. D. (2016). Perceptual learning to discriminate the intensity and spatial location of nociceptive stimuli. Scientific Reports, 6(1), 39104. https://doi.org/10.1038/srep39104

Maris, E., & Oostenveld, R. (2007). Nonparametric statistical testing of EEG- and MEG-data. Journal of Neuroscience Methods, 164(1), 177–190. https://doi.org/10.1016/j.jneumeth.2007.03.024

Mathewson, K. E., Fabiani, M., Gratton, G., Beck, D. M., & Lleras, A. (2010). Rescuing stimuli from invisibility: Inducing a momentary release from visual masking with pre-target entrainment. Cognition, 115(1), 186–191. https://doi.org/10.1016/j.cognition.2009.11.010

Max, C., Widmann, A., Schröger, E., & Sussman, E. (2015). Effects of explicit knowledge and predictability on auditory distraction and target performance. International Journal of Psychophysiology, 98(2), 174–181. https://doi.org/10.1016/j.ijpsycho.2015.09.006

Mento, G., Tarantino, V., Sarlo, M., & Bisiacchi, P. S. (2013). Automatic temporal expectancy: a high-density event-related potential study. PloS One, 8(5), e62896. https://doi.org/10.1371/journal.pone.0062896

Mühlberg, S., Oriolo, G., & Soto-Faraco, S. (2014). Cross-modal decoupling in temporal attention. The European Journal of Neuroscience, 39(12), 2089–2097. https://doi.org/10.1111/ejn.12563

Müller, H. J., & Rabbitt, P. M. (1989). Reflexive and voluntary orienting of visual attention: time course of activation and resistance to interruption. Journal of Experimental Psychology. Human Perception and Performance, 15(2), 315–330. Retrieved from http://www.ncbi.nlm.nih.gov/pubmed/2525601

Nissen, M. J., & Bullemer, P. (1987). Attentional requirements of learning: Evidence from performance measures. Cognitive Psychology, 19(1), 1–32. https://doi.org/10.1016/0010-0285(87)90002-8

Nobre, Anna Christina, & Rohenkohl, G. (2014). Time for the Fourth Dimension in Attention. In A C Nobre & S. Kastner (Eds.), The Oxford Handbook of Attention (pp. 676–724). Oxford University Press.

Noesselt, T., Rieger, J. W., Schoenfeld, M. A., Kanowski, M., Hinrichs, H., Heinze, H.-J., & Driver, J. (2007). Audiovisual temporal correspondence modulates human multisensory superior temporal sulcus plus primary sensory cortices. The Journal of Neuroscience: The Official Journal of the Society for Neuroscience, 27(42), 11431–11441. https://doi.org/10.1523/JNEUROSCI.2252-07.2007

Noesselt, T., Tyll, S., Boehler, C. N., Budinger, E., Heinze, H.-J., & Driver, J. (2010). Sound-induced enhancement of low-intensity vision: multisensory influences on human sensory-specific cortices and thalamic bodies relate to perceptual enhancement of visual detection sensitivity. The Journal of Neuroscience: The Official Journal of the Society for Neuroscience, 30(41), 13609–13623. https://doi.org/10.1523/JNEUROSCI.4524-09.2010

Parise, C. V, Spence, C., & Ernst, M. O. (2012). When correlation implies causation in multisensory integration. Current Biology: CB, 22(1), 46–49. https://doi.org/10.1016/j.cub.2011.11.039

Preston, A. R., & Gabrieli, J. D. E. (2008). Dissociation between Explicit Memory and Configural Memory in the Human Medial Temporal Lobe. Cerebral Cortex, 18(9), 2192–2207. https://doi.org/10.1093/cercor/bhm245

Roach, N. W., Heron, J., & McGraw, P. V. (2006). Resolving multisensory conflict: a strategy for balancing the costs and benefits of audio-visual integration. Proceedings. Biological Sciences, 273(1598), 2159–2168. https://doi.org/10.1098/rspb.2006.3578

Rohenkohl, G., Coull, J. T., & Nobre, A. C. (2011). Behavioural dissociation between exogenous and endogenous temporal orienting of attention. PloS One, 6(1), e14620. https://doi.org/10.1371/journal.pone.0014620

Rohenkohl, G., Cravo, A. M., Wyart, V., & Nobre, A. C. (2012). Temporal expectation improves the quality of sensory information. The Journal of Neuroscience: The Official Journal of the Society for Neuroscience, 32(24), 8424–8428. https://doi.org/10.1523/JNEUROSCI.0804-12.2012

Sanchez, D. J., & Reber, P. J. (2013). Explicit pre-training instruction does not improve implicit perceptual-motor sequence learning. Cognition, 126(3), 341–351. https://doi.org/10.1016/j.cognition.2012.11.006

Sassenhagen, J., & Draschkow, D. (2019). Cluster-based permutation tests of MEG/EEG data do not establish significance of effect latency or location. Psychophysiology, 56(6), e13335. https://doi.org/10.1111/psyp.13335

Schlagbauer, B., Muller, H. J., Zehetleitner, M., & Geyer, T. (2012). Awareness in contextual cueing of visual search as measured with concurrent access- and phenomenal-consciousness tasks. Journal of Vision, 12(11), 25–25. https://doi.org/10.1167/12.11.25

Seitz, A. R. (2017). Perceptual learning. Current Biology, 27(13), R631–R636. https://doi.org/10.1016/j.cub.2017.05.053

Seitz, A. R., & Watanabe, T. (2009). The phenomenon of task-irrelevant perceptual learning. Vision Research, 49(21), 2604–2610. https://doi.org/10.1016/j.visres.2009.08.003

Shen, D., & Alain, C. (2012). Implicit temporal expectation attenuates auditory attentional blink. PloS One, 7(4), e36031. https://doi.org/10.1371/journal.pone.0036031

Smith, A. C., & Brown, E. N. (2003). Estimating a State-Space Model from Point Process Observations. Neural Computation, 15(5), 965–991. https://doi.org/10.1162/089976603765202622

Smith, A. C., Frank, L. M., Wirth, S., Yanike, M., Hu, D., Kubota, Y., … Brown, E. N. (2004). Dynamic analysis of learning in behavioral experiments. The Journal of Neuroscience: The Official Journal of the Society for Neuroscience, 24(2), 447–461. https://doi.org/10.1523/JNEUROSCI.2908-03.2004

Starke, J., Ball, F., Heinze, H.-J., & Noesselt, T. (2017). The spatio-temporal profile of multisensory integration. European Journal of Neuroscience. https://doi.org/10.1111/ejn.13753 [EPub ahead of print]

Stefaniak, N., Willems, S., Adam, S., & Meulemans, T. (2008). What is the impact of the explicit knowledge of sequence regularities on both deterministic and probabilistic serial reaction time task performance? Memory & Cognition, 36(7), 1283–1298. https://doi.org/10.3758/MC.36.7.1283

Summerfield, C., & Egner, T. (2009). Expectation (and attention) in visual cognition. Trends in Cognitive Sciences, 13(9), 403–409. https://doi.org/10.1016/j.tics.2009.06.003

Taylor, J. A., & Ivry, R. B. (2013). Implicit and Explicit Processes in Motor Learning. In Action Science (pp. 63–87). The MIT Press. https://doi.org/10.7551/mitpress/9780262018555.003.0003

Turk-Browne, N. B., Scholl, B. J., Chun, M. M., & Johnson, M. K. (2009). Neural Evidence of Statistical Learning: Efficient Detection of Visual Regularities Without Awareness. Journal of Cognitive Neuroscience, 21(10), 1934–1945. https://doi.org/10.1162/jocn.2009.21131

Turk-Browne, N. B., Scholl, B. J., Johnson, M. K., & Chun, M. M. (2010). Implicit perceptual anticipation triggered by statistical learning. The Journal of Neuroscience: The Official Journal of the Society for Neuroscience, 30(33), 11177–11187. https://doi.org/10.1523/JNEUROSCI.0858-10.2010

Vadillo, M. A., Konstantinidis, E., & Shanks, D. R. (2016). Underpowered samples, false negatives, and unconscious learning. Psychonomic Bulletin & Review, 23(1), 87–102. https://doi.org/10.3758/s13423-015-0892-6

Van den Bussche, E., Vermeiren, A., Desender, K., Gevers, W., Hughes, G., Verguts, T., & Reynvoet, B. (2013). Disentangling conscious and unconscious processing: a subjective trial-based assessment approach. Frontiers in Human Neuroscience, 7, 769. https://doi.org/10.3389/fnhum.2013.00769

Vatakis, A., Bayliss, L., Zampini, M., & Spence, C. (2007). The influence of synchronous audiovisual distractors on audiovisual temporal order judgments. Perception & Psychophysics, 69(2), 298–309. https://doi.org/10.3758/bf03193751

Wagenmakers, E.-J., & Farrell, S. (2004). AIC model selection using Akaike weights. Psychonomic Bulletin & Review, 11(1), 192–196. Retrieved from http://www.ncbi.nlm.nih.gov/pubmed/15117008

Wang, P., & Nikolić, D. (2011). An LCD Monitor with Sufficiently Precise Timing for Research in Vision. Frontiers in Human Neuroscience, 5(85), 85. https://doi.org/10.3389/fnhum.2011.00085

Warner, C. B., Juola, J. F., & Koshino, H. (1990). Voluntary allocation versus automatic capture of visual attention. Perception & Psychophysics, 48(3), 243–251. https://doi.org/10.3758/BF03211524

Werner, S., & Noppeney, U. (2010). Superadditive Responses in Superior Temporal Sulcus Predict Audiovisual Benefits in Object Categorization. Cerebral Cortex, 20(8), 1829–1842. https://doi.org/10.1093/cercor/bhp248

Westerberg, C. E., Miller, B. B., Reber, P. J., Cohen, N. J., & Paller, K. A. (2011). Neural correlates of contextual cueing are modulated by explicit learning. Neuropsychologia, 49(12), 3439–3447. https://doi.org/10.1016/j.neuropsychologia.2011.08.019

Wirth, S., Yanike, M., Frank, L. M., Smith, A. C., Brown, E. N., & Suzuki, W. A. (2003). Single Neurons in the Monkey Hippocampus and Learning of New Associations. Science, 300(5625), 1578–1581. https://doi.org/10.1126/science.1084324

Zanto, T. P., Pan, P., Liu, H., Bollinger, J., Nobre, A. C., & Gazzaley, A. (2011). Age-related changes in orienting attention in time. The Journal of Neuroscience: The Official Journal of the Society for Neuroscience, 31(35), 12461–12470. https://doi.org/10.1523/JNEUROSCI.1149-11.2011

