## Supplement for "Minimal interplay between explicit knowledge, dynamics of learning and temporal expectations in different, complex uni- and multisensory contexts"

### Supplement 1: Re-analysis Ball et al. (2018)

*Description: Given that we do not exclude, but transform outliers in the current study (conversion of trials with < 150 ms RT to incorrect trials), we re-analysed all experimental data using this criterion to check whether it would change any results. As in Ball et al. (2018) mean accuracies were converted to d-prime and ANOVAs were conducted for each experiment and performance measure (A1-4: d-prime, B1-4: RT). The results are virtually identical with our previous report. Note that we also show plots of the TE*Modality data (with 95% confidence intervals as calculated in JASP). Abbreviations: df = degrees of freedom, F = F-statistic, t = t-statistic, p/pbonf = p value/p value Bonferroni corrected, η² = eta-squared, TE = temporal expectation, AV = audio-visual, A = auditory, V = visual, C1/C2 = contrasted conditions in post hoc test.*

#### *A1: Experiment 1 - rm ANOVA (accuracy scores)*

| effect | Sphericity Correction | df | F | p | η² |
|---|---|---|---|---|---|
| TE | ----- | 1 | 28.291 | < .001 | 0.043 |
| Residual | ----- | 29 |  |  |  |
| Modality | GG | 1.297 | 2.495 | 0.115 | 0.065 |
| Residual | GG | 37.627 | | | |
| TE ⁎ Modality | GG | 1.659 | 0.655 | 0.496 | 0.002 |
| Residual | GG | 48.118 | | | |

*Post hoc comparison for main effect TE and main effect Modality (post hoc effects in JASP)*

| C1 | C2 | Mean Difference | SE | t | $p_{bonf}$ |
|---|---|---|---|---|---|
| Unexpected | Expected | -0.167 | 0.031 | -5.319 | < .001 |
| AV | A | 0.246 | 0.113 | 2.183 | 0.099 |
| | V | 0.169 | 0.113 | 1.501 | 0.417 |
| A | V | -0.077 | 0.113 | -0.683 | 1 |

*Data plot TE * Modality (individual curves are expected/unexpected trials)*

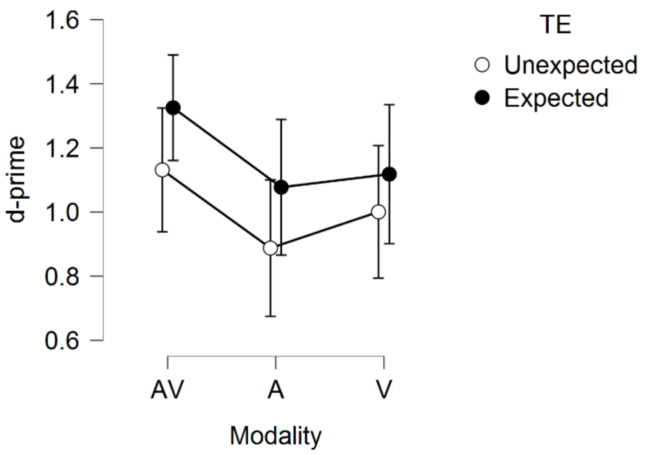

***A2: Experiment 2 - rm ANOVA (accuracy scores))***

| effect | Sphericity Correction | df | F | p | $\eta^2$ |
|---|---|---|---|---|---|
| TE | ----- | 1 | 27.286 | < .001 | 0.078 |
| Residual | ----- | 29 |  |  |  |
| Modality | GG | 1.467 | 9.14 | 0.001 | 0.166 |
| Residual | GG | 42.55 | | | |
| TE ∗ Modality | ----- | 2 | 6.705 | 0.002 | 0.027 |
| Residual | ----- | 58 |  |  |  |

*Post hoc comparison for main effect TE and main effect Modality (post hoc effects in JASP)*

| C1 | C2 | Mean Difference | SE | t | $p_{bonf}$ |
|---|---|---|---|---|---|
| Unexpected | Expected | -0.245 | 0.047 | -5.224 | < .001 |
| AV | A | 0.308 | 0.102 | 3.022 | 0.011 |
| | V | 0.421 | 0.102 | 4.13 | < .001 |
| A | V | 0.113 | 0.102 | 1.109 | 0.816 |

*Post hoc comparison for factor TE (expected vs. unexpected) for each modality type (simple main effects in JASP)*

| Modality | df | F | p |
|---|---|---|---|
| AV | 1 | 24.14 | < .001 |
| A | 1 | 14.593 | < .001 |
| V | 1 | 0.63 | 0.434 |

*Data plot TE * Modality (individual curves are expected/unexpected trials)*

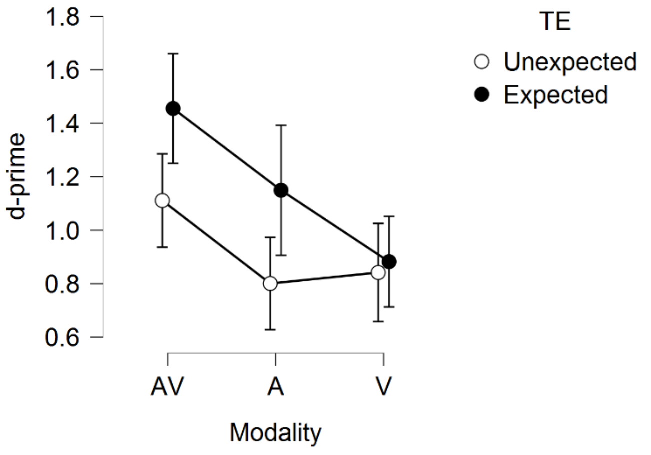

### *A3: Experiment 3 - rm ANOVA (accuracy scores)*

| effect | Sphericity Correction | df | F | p | η² |
|---|---|---|---|---|---|
| TE | ----- | 1 | 19.721 | < .001 | 0.027 |
| Residual | ----- | 29 |  |  |  |
| Modality | GG | 1.339 | 17.564 | < .001 | 0.317 |
| Residual | GG | 38.839 | | | |
| TE * Modality | ----- | 2 | 0.157 | 0.855 | 5.087e -4 |
| Residual | ----- | 58 |  |  |  |

*Post hoc comparison for main effect TE and main effect Modality (post hoc effects in JASP)*

| C1 | C2 | Mean Difference | SE | t | $p_{bonf}$ |
|---|---|---|---|---|---|
| Unexpected | Expected | -0.143 | 0.032 | -4.441 | < .001 |
| AV | A | 0.522 | 0.102 | 5.144 | < .001 |
| | V | 0.52 | 0.102 | 5.122 | < .001 |
| A | V | -0.002 | 0.102 | -0.021 | 1 |

*Data plot TE * Modality (individual curves are expected/unexpected trials)*

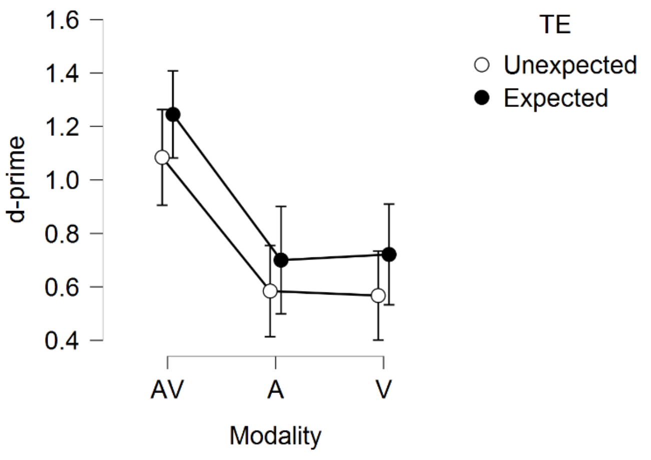

#### *A4: Experiment 4 - rm ANOVA (accuracy scores)*

| effect | Sphericity Correction | df | F | p | $\eta^2$ |
|---|---|---|---|---|---|
| TE | ----- | 1 | 7.238 | 0.012 | 0.012 |
| Residual | ----- | 29 |  |  |  |
| Modality | GG | 1.551 | 18.843 | < .001 | 0.34 |
| Residual | GG | 44.985 | | | |
| TE ∗ Modality | ----- | 2 | 3.628 | 0.033 | 0.009 |
| Residual | ----- | 58 |  |  |  |

*Post hoc comparison for main effect TE and main effect Modality (post hoc effects in JASP)*

| C1 | C2 | Mean Difference | SE | t | $p_{bonf}$ |
|---|---|---|---|---|---|
| Unexpected | Expected | -0.093 | 0.034 | -2.69 | 0.012 |
| AV | A | 0.489 | 0.099 | 4.935 | < .001 |
| | V | 0.558 | 0.099 | 5.629 | < .001 |
| A | V | 0.069 | 0.099 | 0.694 | 1 |

*Post hoc comparison for factor TE (expected vs. unexpected) for each modality type (simple main effects in JASP)*

| Modality | df | F | p |
|---|---|---|---|
| AV | 1 | 12.043 | 0.002 |
| A | 1 | 2.179 | 0.151 |
| V | 1 | 0.034 | 0.856 |

*Data plot TE * Modality (individual curves are expected/unexpected trials)*

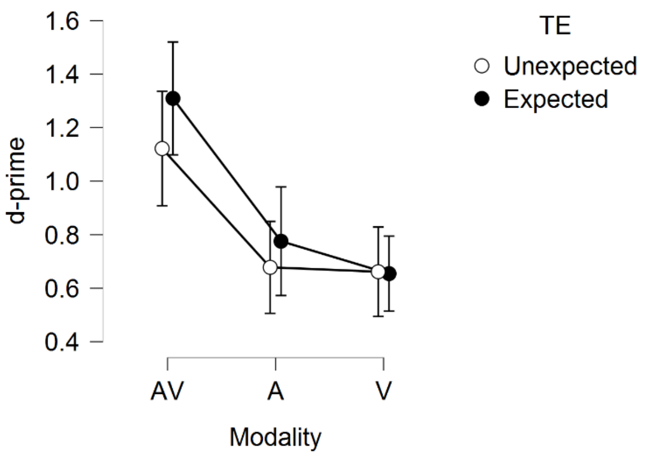

**_B1: Experiment 1 - rm ANOVA (RT scores)_**

| effect | Sphericity Correction | df | F | p | η² |
|---|---|---|---|---|---|
| TE | ----- | 1 | 33.453 | < .001 | 0.091 |
| Residual | ----- | 29 |  |  |  |
| Modality | GG | 1.335 | 4.011 | 0.041 | 0.094 |
| Residual | GG | 38.718 | | | |
| TE ∗ Modality | ----- | 2 | 5.13 | 0.009 | 0.009 |
| Residual | ----- | 58 |  |  |  |

_Post hoc comparison for main effect TE and main effect Modality (post hoc effects in JASP)_

| C1 | C2 | Mean Difference | SE | t | $p_{bonf}$ |
|---|---|---|---|---|---|
| Unexpected | Expected | 0.127 | 0.022 | 5.784 | < .001 |
| AV | A | -0.143 | 0.056 | -2.546 | 0.041 |
| | V | -0.131 | 0.056 | -2.347 | 0.067 |
| A | V | 0.011 | 0.056 | 0.199 | 1 |

_Post hoc comparison for factor TE (expected vs. unexpected) for each modality type (simple main effects in JASP)_

| Modality | df | F | p |
|---|---|---|---|
| AV | 1 | 26.026 | < .001 |
| A | 1 | 37.042 | < .001 |
| V | 1 | 6.569 | 0.016 |

_Data plot TE * Modality (individual curves are expected/unexpected trials)_

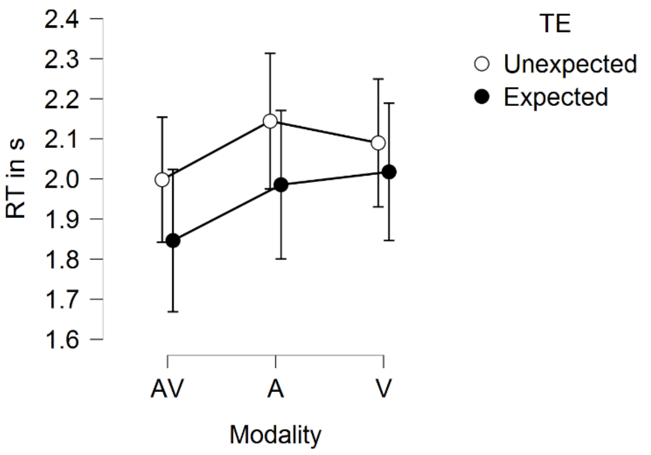

### *B2: Experiment 2 - rm ANOVA (RT scores)*

| effect | Sphericity Correction | df | F | p | $\eta^2$ |
|---|---|---|---|---|---|
| TE | ----- | 1 | 20.999 | < .001 | 0.163 |
| Residual | ----- | 29 |  |  |  |
| Modality | GG | 1.254 | 5.587 | 0.017 | 0.087 |
| Residual | GG | 36.354 | | | |
| TE ∗ Modality | ----- | 2 | 5.794 | 0.005 | 0.013 |
| Residual | ----- | 58 |  |  |  |

*Post hoc comparison for main effect TE and main effect Modality (post hoc effects in JASP)*

| C1 | C2 | Mean Difference | SE | t | $p_{bonf}$ |
|---|---|---|---|---|---|
| Unexpected | Expected | 0.142 | 0.031 | 4.582 | < .001 |
| AV | A | -0.1 | 0.038 | -2.621 | 0.034 |
| | V | -0.118 | 0.038 | -3.107 | 0.009 |
| A | V | -0.018 | 0.038 | -0.486 | 1 |

*Post hoc comparison for factor TE (expected vs. unexpected) for each modality type (simple main effects in JASP)*

| Modality | df | F | p |
|---|---|---|---|
| AV | 1 | 16.63 | < .001 |
| A | 1 | 20.51 | < .001 |
| V | 1 | 12.82 | 0.001 |

*Data plot TE * Modality (individual curves are expected/unexpected trials)*

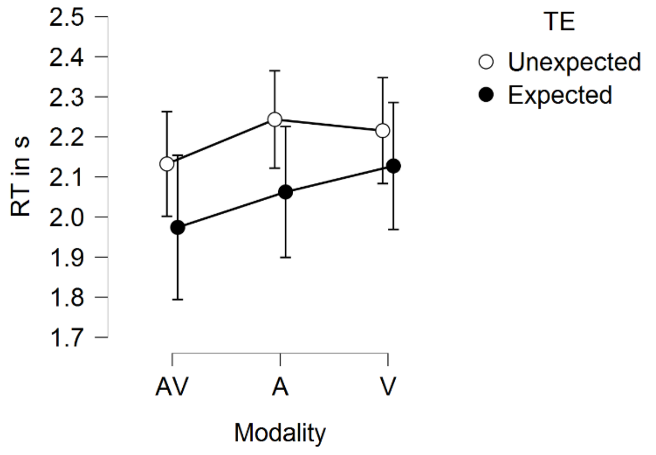

***B3: Experiment 3 - rm ANOVA (RT scores)***

| effect | Sphericity Correction | df | F | p | η² |
|---|---|---|---|---|---|
| TE | ----- | 1 | 18.272 | < .001 | 0.052 |
| Residual | ----- | 29 |  |  |  |
| Modality | GG | 1.367 | 20.232 | < .001 | 0.342 |
| Residual | GG | 39.63 | | | |
| TE ⁎ Modality | ----- | 2 | 1.366 | 0.263 | 0.001 |
| Residual | ----- | 58 |  |  |  |

*Post hoc comparison for main effect TE and main effect Modality (post hoc effects in JASP)*

| C1 | C2 | Mean Difference | SE | t | $p_{bonf}$ |
|---|---|---|---|---|---|
| Unexpected | Expected | 0.098 | 0.023 | 4.275 | < .001 |
| AV | A | -0.208 | 0.048 | -4.307 | < .001 |
| | V | -0.3 | 0.048 | -6.207 | < .001 |
| A | V | -0.092 | 0.048 | -1.901 | 0.187 |

*Data plot TE * Modality (individual curves are expected/unexpected trials)*

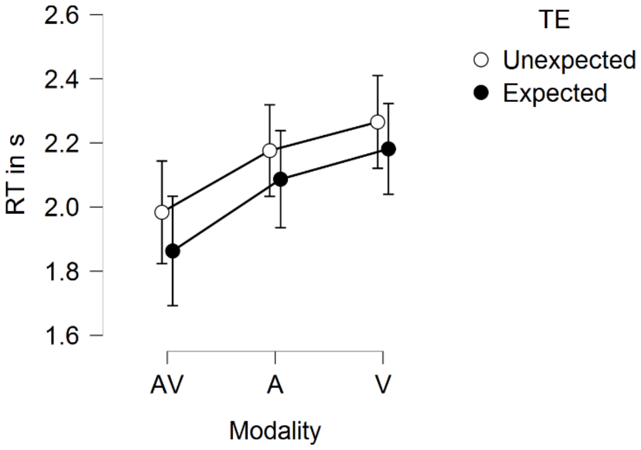

#### *B4: Experiment 4 - rm ANOVA (RT scores)*

| **effect** | **Sphericity Correction** | **df** | **F** | **p** | **η²** |
|---|---|---|---|---|---|
| TE | ----- | 1 | 15.156 | < .001 | 0.047 |
| Residual | ----- | 29 |  |  |  |
| Modality | GG | 1.507 | 17.253 | < .001 | 0.304 |
| Residual | GG | 43.717 | | | |
| TE ⁎ Modality | ----- | 2 | 1.015 | 0.369 | 0.002 |
| Residual | ----- | 58 |  |  |  |

*Post hoc comparison for main effect TE and main effect Modality (post hoc effects in JASP)*

| **C1** | **C2** | **Mean Difference** | **SE** | **t** | **p bonf** |
|---|---|---|---|---|---|
| Unexpected | Expected | 0.082 | 0.021 | 3.893 | < .001 |
| AV | A | -0.206 | 0.044 | -4.721 | < .001 |
| | V | -0.235 | 0.044 | -5.388 | < .001 |
| A | V | -0.029 | 0.044 | -0.667 | 1 |

*Data plot TE * Modality (individual curves are expected/unexpected trials)*

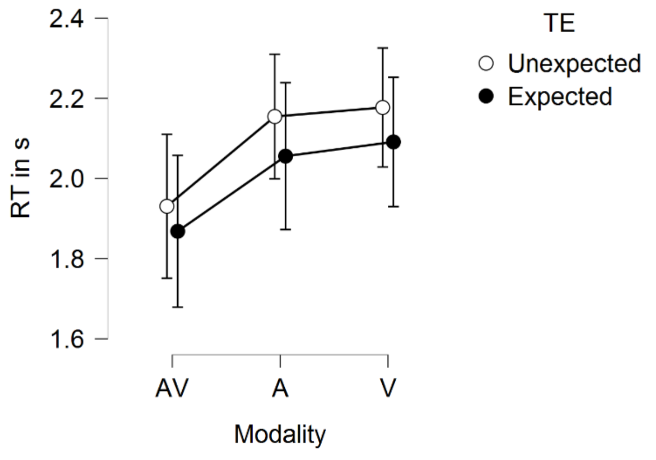

### Supplement 2: Mixed Model (MM) design in R

*Description: R code for computing models for different performance measures.*

*A1/2: models for accuracy and RT (split for whole data set and modality-specific analyses)*
*B1/2: models for learning trials (split for whole data set and modality-specific analyses)*

*Perf: accuracy, RT oder learn trial scores*
*TE: Temporal Expectation - factor with 2 levels (1 = no, 2 = yes)*
*Mod: Modality - factor with 3 levels (1 = AV, 2 = A, 3 = V)*
*SpatUnc: Spatial Uncertainty - factor with 2 levels (1 = low, 2 = high)*
*Know: Knowledge group - factor with 2 levels (1 = implicit, 2 = explicit)*
*Run: Run (which part of experiment) - factor with 3 levels (1 = run1/2, 2 = run3/4, 3 = run5/6)*
*MSUnc : Modality-specific Uncertainty - factor with 2 levels (1 = low, 2 = high)*

*Each model was run with: options(contrasts = c("contr.sum","contr.poly"))*

#### *A1: MM for accuracy & RT data (InputData = whole data set)*

```
TM1 <- mixed(Perf ~ 1 + (1|MSUnc) + (1|id), data = InputData, method="KR")
TM2 <- mixed(Perf ~ 1 + TE + (1|MSUnc) + (1|id), data = InputData, method="KR")
TM3 <- mixed(Perf ~ 1 + TE*Mod + (1|MSUnc) + (1|id), data = InputData, method="KR")
TM4 <- mixed(Perf ~ 1 + TE*SpatUnc + (1|MSUnc) + (1|id), data = InputData, method="KR")
TM5 <- mixed(Perf ~ 1 + TE*Know + (1|MSUnc) + (1|id), data = InputData, method="KR")
TM6 <- mixed(Perf ~ 1 + TE*Run + (1|MSUnc) + (1|id), data = InputData, method="KR")
TM7 <- mixed(Perf ~ 1 + TE*Mod*SpatUnc + (1|MSUnc) + (1|id), data = InputData, method="KR")
TM8 <- mixed(Perf ~ 1 + TE*Mod*Know + (1|MSUnc) + (1|id), data = InputData, method="KR")
TM9 <- mixed(Perf ~ 1 + TE*Mod*Run + (1|MSUnc) + (1|id), data = InputData, method="KR")
TM10 <- mixed(Perf ~ 1 + TE*SpatUnc*Know + (1|MSUnc) + (1|id), data = InputData, method="KR")
TM11 <- mixed(Perf ~ 1 + TE*SpatUnc*Run + (1|MSUnc) + (1|id), data = InputData, method="KR")
TM12 <- mixed(Perf ~ 1 + TE*Know*Run + (1|MSUnc) + (1|id), data = InputData, method="KR")
TM13 <- mixed(Perf ~ 1 + TE*Mod*SpatUnc*Know + (1|MSUnc) + (1|id), data = InputData, method="KR")
TM14 <- mixed(Perf ~ 1 + TE*Mod*SpatUnc*Run + (1|MSUnc) + (1|id), data = InputData, method="KR")
TM15 <- mixed(Perf ~ 1 + TE*Mod*Know*Run + (1|MSUnc) + (1|id), data = InputData, method="KR")
TM16 <- mixed(Perf ~ 1 + TE*Mod*SpatUnc*Run*Know + (1|MSUnc) + (1|id), data = InputData, method="KR")
```

#### *A2: MM for accuracy & RT data (InputData = low OR high modality-spec. uncertainty)*

```
TM1 <- mixed(Perf ~ 1 + (1|id), data = InputData, method="KR")
TM2 <- mixed(Perf ~ 1 + TE + (1|id), data = InputData, method="KR")
TM3 <- mixed(Perf ~ 1 + TE*Mod + (1|id), data = InputData, method="KR")
TM4 <- mixed(Perf ~ 1 + TE*SpatUnc + (1|id), data = InputData, method="KR")
TM5 <- mixed(Perf ~ 1 + TE*Know + (1|id), data = InputData, method="KR")
TM6 <- mixed(Perf ~ 1 + TE*Run + (1|id), data = InputData, method="KR")
TM7 <- mixed(Perf ~ 1 + TE*Mod*SpatUnc + (1|id), data = InputData, method="KR")
TM8 <- mixed(Perf ~ 1 + TE*Mod*Know + (1|id), data = InputData, method="KR")
TM9 <- mixed(Perf ~ 1 + TE*Mod*Run + (1|id), data = InputData, method="KR")
TM10 <- mixed(Perf ~ 1 + TE*SpatUnc*Know + (1|id), data = InputData, method="KR")
TM11 <- mixed(Perf ~ 1 + TE*SpatUnc*Run + (1|id), data = InputData, method="KR")
TM12 <- mixed(Perf ~ 1 + TE*Know*Run + (1|id), data = InputData, method="KR")
TM13 <- mixed(Perf ~ 1 + TE*Mod*SpatUnc*Know + (1|id), data = InputData, method="KR")
TM14 <- mixed(Perf ~ 1 + TE*Mod*SpatUnc*Run + (1|id), data = InputData, method="KR")
TM15 <- mixed(Perf ~ 1 + TE*Mod*Know*Run + (1|id), data = InputData, method="KR")
TM16 <- mixed(Perf ~ 1 + TE*Mod*SpatUnc*Run*Know + (1|id), data = InputData, method="KR")
```

### *B1: MM for learn trial data (InputData = whole data set)*

```
LTM1  <- mixed(Perf ~ 1 + (1|MSUnc) + (1|id), data = InputData, method="KR")
LTM2  <- mixed(Perf ~ 1 + Mod + (1|MSUnc) + (1|id), data = InputData, method="KR")
LTM3  <- mixed(Perf ~ 1 + Know + (1|MSUnc) + (1|id), data = InputData, method="KR")
LTM4 <- mixed(Perf ~ 1 + SpatUnc + (1|MSUnc) + (1|id), data = InputData, method="KR")
LTM5  <- mixed(Perf ~ 1 + Mod*Know + (1|MSUnc) + (1|id), data = InputData, method="KR")
LTM6  <- mixed(Perf ~ 1 + Mod*SpatUnc + (1|MSUnc) + (1|id), data = InputData, method="KR")
LTM7  <- mixed(Perf ~ 1 + SpatUnc*Know + (1|MSUnc) + (1|id), data = InputData, method="KR")
LTM8  <- mixed(Perf ~ 1 + Mod*SpatUnc*Know + (1|MSUnc) + (1|id), data = InputData, method="KR")
```

### *B2: MM for learn trial data (InputData = low OR high modality-spec. uncertainty)*

```
LTM1  <- mixed(Perf ~ 1 + (1|id), data = InputData, method="KR")
LTM2  <- mixed(Perf ~ 1 + Mod + (1|id), data = InputData, method="KR")
LTM3  <- mixed(Perf ~ 1 + Know + (1|id), data = InputData, method="KR")
LTM4 <- mixed(Perf ~ 1 + SpatUnc + (1|id), data = InputData, method="KR")
LTM5  <- mixed(Perf ~ 1 + Mod*Know + (1|id), data = InputData, method="KR")
LTM6  <- mixed(Perf ~ 1 + Mod*SpatUnc + (1|id), data = InputData, method="KR")
LTM7  <- mixed(Perf ~ 1 + SpatUnc*Know + (1|id), data = InputData, method="KR")
LTM8  <- mixed(Perf ~ 1 + Mod*SpatUnc*Know + (1|id), data = InputData, method="KR")
```
